# Chemogenetic activation of target neurons expressing insect Ionotropic Receptors in the mammalian central nervous system by systemic administration of ligand precursors

**DOI:** 10.1101/2023.09.11.557081

**Authors:** Yoshio Iguchi, Ryoji Fukabori, Shigeki Kato, Kazumi Takahashi, Satoshi Eifuku, Yuko Maejima, Kenju Shimomura, Hiroshi Mizuma, Aya Mawatari, Hisashi Yilong Cui, Hirotaka Onoe, Keigo Hikishima, Makoto Osanai, Takuma Nishijo, Toshihiko Momiyama, Richard Benton, Kazuto Kobayashi

**Affiliations:** Department of Molecular Genetics, Institute of Biomedical Sciences, Fukushima Medical University School of Medicine, Fukushima, Japan; Department of Systems Neuroscience, Fukushima Medical University School of Medicine, Fukushima Japan; Department of Bioregulation and Pharmacological Medicine, Fukushima Medical University School of Medicine, Fukushima Japan; Laboratory for Pathophysiological and Health Science, RIKEN Center for Biosystems Dynamics Research, Kobe, Japan; Department of Functional Brain Imaging, Institute for Quantum Medical Science, National Institutes for Quantum Science and Technology, Chiba, Japan; Laboratory for Labeling Chemistry, RIKEN Center for Biosystems Dynamics Research, Kobe, Japan; Research Institute for Drug Discovery Science, Collaborative Creation Research Center, Organization for Research Promotion, Osaka Metropolitan University; Laboratory for Biofunction Dynamics Imaging, RIKEN Center for Biosystems Dynamics Research, Kobe, Japan; Human Brain Research Center, Kyoto University Graduate School of Medicine, Kyoto, Japan; Medical Devices Research Group, Health and Medical Research Institute, National Institute of Advanced Industrial Science and Technology (AIST), Tsukuba, Japan; Department of Medical Physics and Engineering, Division of Health Sciences, Osaka University Graduate School of Medicine, Suita, Japan; Department of Pharmacology, Jikei University School of Medicine, Tokyo, Japan; Department of Molecular Neurobiology, Institute for Developmental Research, Aichi Developmental Disability Center, Kasugai, Japan; Center for Integrative Genomics, Faculty of Biology and Medicine, University of Lausanne, Lausanne, Switzerland

## Abstract

The IR-mediated neuronal activation (IRNA) technology allows stimulation of neurons in the brain that heterologously-express members of the insect chemosensory IR repertoire in response to their cognate ligands. In the current protocol, a ligand against the complex consisting of IR84a and IR8a subunits, phenylacetic acid (PhAc), is locally injected into a brain region, because of a low efficiency of PhAc for the delivery into the brain across the blood-brain barrier. To circumvent this invasive injection, here we developed a strategy for activation of target neurons in the brain through peripheral administration with a precursor of PhAc, methyl ester of PhAc (PhAcM), which is efficiently transferred into the brain and converted to the mature ligand by endogenous esterase activities. Peripheral administration of with PhAcM activated IR84a/IR8a-expressing neurons in the locus coeruleus of mice and increased the release of neurotransmitters in their nerve terminal regions. (S)-2-phenylpropionic acid ((S)-PhPr) was newly identified as a ligand for IR84a/IR8a, and peripheral administration with the methyl ester of PhPr with the S-configuration [(S)-PhPrM] caused similar effects on the target neurons. In addition, cell-type specific expression of IR84a/IR8a complex in the striatum of rats was unilaterally induced with a viral vector based on the Cre-loxP system. Peripheral administration with PhAcM or (S)-PhPrM stimulated the neurotransmitter release in the ipsilateral terminal regions of the vector-injected striatum, and PhAcM administration resulted in rotational behavior.

Finally, we demonstrated that the metabolites of the peripherally administered-radiolabeled (*S*)-PhPrM accumulated in the IR84a/IR8a-expressing region in the striatum of the vector-injected rats. These results demonstrate that the systemic IRNA technique provides a powerful strategy for remote manipulation of diverse types of target neurons in the mammalian central nervous system.

## Introduction

Uncovering the roles of specific neuronal populations and neural pathways constituting the complex central nervous system is essential for understanding the mechanisms of higher brain function and the pathophysiology of various psychiatric and neurological disorders. Sophisticated genetic approaches, including transgenic animals and viral vector systems, have provided a powerful experimental strategy for manipulating the functions of these neuronal types and pathways (Carter, 1993; Kobayashi et al., 2018). Among them, an increasingly important approach is the chemogenetic manipulation of target neuronal populations (Atasoy and Sternson, 2018; Sternson and Roth, 2014). A common class of chemogenetic tool is DREADDs, which are genetically engineered G protein-coupled receptors exclusively activated by designer drugs (Armbruster et al., 2007; Roth, 2016). DREADDs have made notable progress, including a full line-up of engineered receptors (Nakajima and Wess, 2012; Vardy et al., 2015) and the development of ligands with high affinity to the receptor but minor off-target effects (Chen et al., 2015; Nagai et al., 2020). However, they work through altered intracellular signaling of target neurons, which might induce complex cellular reactions (Van Steenbergen and Bareyre, 2021). For example, DREADD hM3D (Gq)-dependent pre– and postsynaptic mechanisms exhibited different ligand dose responses (Pati et al., 2019). Furthermore, a prolonged high-level DREADD hM4D (Gi) expression resulted in neurotoxic effects (Goossens et al., 2021).

Another chemogenetic manipulation approach is using ligand-gated ion channels (LGICs) that do not require intracellular signaling pathways to modify neuronal activity (Atasoy and Sternson, 2018; Magnus et al., 2019). LGICs are generally classified into the Cys-loop, P2X, and ionotropic glutamate receptor (iGluR) families. Among them, several approaches depending on Cys-loop receptors have been studied, such as silencing of neuronal activity using glutamate/ivermectin-gated chloride channels derived from *C. elegans* (Lerchner et al., 2007; Lin et al., 2011), activation of neuron using an ivermectin-sensitive glycine receptor mutant with cation permeability (Islam et al., 2016), and a system using chimeric receptors in which the ligand binding domains of Cys-loop ion channels (pharmacologically selective actuator modules) activated by cognate synthetic ligands (pharmacologically selective effector molecules) (Magnus et al., 2011; Magnus et al., 2019). Although an ATP-sensitive P2X2-based tool has been reported in invertebrates (Zemelman et al., 2003), knockout mutant backgrounds are required for application in mammals (Atasoy and Sternson, 2018).

Recently, we have developed a new type of chemogenetic tool, the Ionotropic Receptors (IR)-mediated neuronal activation (IRNA) system (Fukabori et al., 2020). Insect IRs belong to the iGluR superfamily and are involved in the sensation of a wide diversity of volatile and non-volatile chemicals as well as in the perception of temperature and humidity (Benton et al., 2009; Rytz et al., 2013; van Giesen and Garrity, 2017). In particular, we used the heteromer of IR84a and IR8a subunits derived from the fly, *Drosophila melanogaster*, which is form a cation channel gated by phenylacetic acid (PhAc) and phenylacetaldehyde (PhAl) (Abuin et al., 2011; Grosjean et al., 2011), to induce activation of target neurons in the mammalian brain. The IR84a/IR8a complex was expressed under the control of the promoter for the gene encoding tyrosine hydroxylase (TH) in transgenic (Tg) mice (TH-IR84a/IR8a), and transgene expression was observed in catecholamine-containing neurons, including norepinephrine (NE) neurons in the locus coeruleus (LC) (Fukabori et al., 2020). In a brain slice preparation, the LC-NE neurons in the Tg mice exhibited an increase in membrane potential and firing frequency in response to PhAc/PhAl. Intracranial microinjection of PhAc into the LC activated NE neurons and elevated the extracellular NE level in the terminal regions, resulting in enhancement of the recall process of emotional memory. These findings indicated the efficacy of IRNA system in activating mammalian target neurons (Fukabori et al., 2020).

In the present study, we developed a strategy of systemic drug administration for the IRNA system to activate target neurons in the brain. The delivery of IR ligands into the brain by systemic administration is less invasive than the intracranial surgery for ligand injection and permits remote activation of the target neurons with minimal disturbance of the animals, particularly in the case of repeated manipulation (Bang et al., 2016; Raper & Galvan., 2022). Since PhAc is known to have the difficulty in crossing the blood-brain barrier (BBB), we used methyl ester of PhAc (PhAcM) for systemic administration, which exhibits lipophilic properties resulting in the facilitated BBB permeability by passive diffusion, and is converted to the mature ligand by esterase activities in the brain. We first performed peripheral administration with PhAcM to activate LC-NE neurons in Tg mice. We then applied this strategy for selective activation of striatal projection neurons containing dopamine (DA) type 2 receptors in the brain of a Cre transgenic rat line (*Drd2*-Cre), in which IR84a/IR8a transgenes were expressed by using a viral vector carrying the flip-excision switch (FLEX) system. Activation of the target neurons by systemic administration of the ligand precursors was confirmed by the elevation of GABA release in their terminal regions and the induction of rotational behavior. Furthermore, we demonstrated the binding of processed ligands in the brain regions expressing the receptor complex using *in vivo* and *ex vivo* brain imaging techniques.

## Results

### Peripheral administration of methyl ester of PhAc induces excitation of the central IR84a/IR8a-expressing neurons

Transfer of PhAc through the BBB is generally considered low efficiency (Pedemonte et al., 1976). To test whether a peripherally administered PhAc can stimulate the target neurons expressing IR84a and IR8a in the brain, we injected PhAc solution intravenously into the lateral tail vein of the TH-IR84a/IR8a mice and monitored NE release in the anterior cingulate cortex (ACC), a brain region innervated by the LC, by using a microdialysis procedure. When we first tested the peripheral administration of PhAc (10 or 30 mg/kg), the extracellular NE level (NE_[ext]_) in the ACC did not show any significant changes as compared to vehicle administration (Figure 1—figure supplement 1A). In contrast, perfusion of different concentrations of PhAc into the dialysis probe (i.e., a reverse dialysis with PhAc) elevated NE_[ext]_ in a dose-dependent manner (Figure 1—figure supplement 1B). These data indicate that the central IR84a/IR8a-expressing neurons are activated by intracranially administered PhAc but respond little to peripherally administered PhAc.

**Figure 1.**
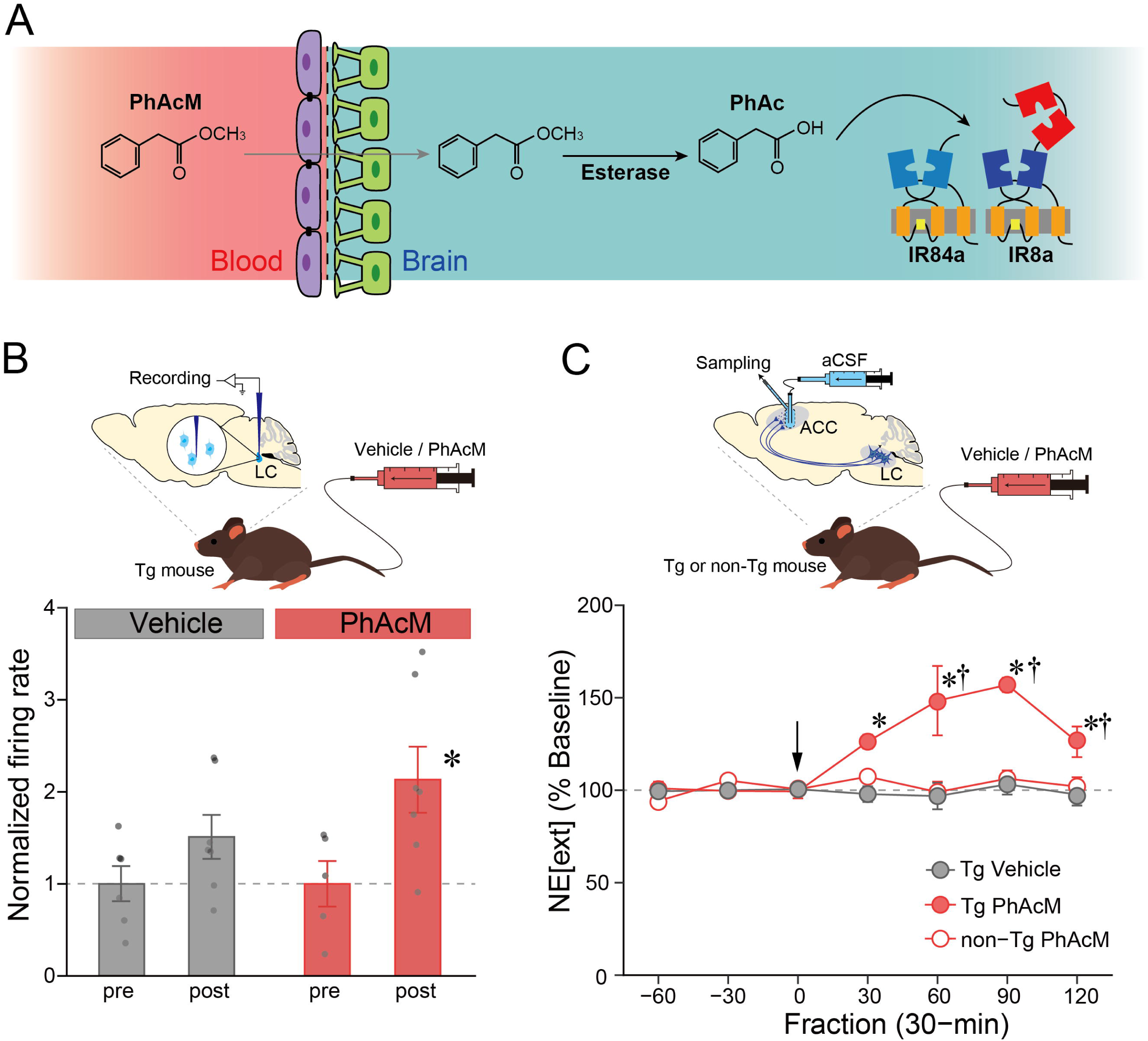
Enhancement of the BBB permeability of PhAc by methyl esterification. (**A**) Schematic representation of drug delivery. PhAcM in the circulating blood is translocated across the BBB and enters into the brain, where PhAcM is converted to PhAc by endogenous esterase activity and PhAc activates the IR84a/IR8a complex. (**B**) Plot of changes in the normalized firing rate after the tail vein injection of PhAcM (20 mg/kg) or vehicle in the TH-IR84a/IR8a (Tg) mice (n = 6 for vehicle/pre, n = 7 for vehicle/post, n = 5 for PhAcM/pre, and n = 7 for PhAcM/post). *p < 0.05 vs. PhAcM/pre. Individual data points are overlaid. (**C**) Plot of changes in NE_[ext]_ in the ACC of Tg and non-transgenic (non-Tg) littermates before and after tail vein injection of PhAcM (10 mg/kg) (n = 4 for each group). NE_[ext]_ is expressed as a percentage of the average baseline levels of each mouse. Arrow indicates the timing of drug injection. NE_[ext]_ in PhAcM-administered Tg mice was significantly higher than that of vehicle-administered Tg mice at the 30– to 120-min fractions (ts_(63)_ = 3.38, 6.12, 6.43, 3.41, p = 0.004, p < 0.001, p < 0.001, p = 0.003, rs = 0.39, 0.61, 0.63, 0.39, respectively) and that of PhAcM-administered non-transgenic (non-Tg) littermates at the 60– to 120-min fractions (ts_(63)_ = 5.90, 6.06, 2.89, p < 0.001, p < 0.001, p = 0.016, rs = 0.60, 0.61, 0.34, respectively). *p < 0.05 vs. Tg/Vehicle; ^†^p < 0.05 vs. non-Tg/PhAcM. Data are presented as mean ± SEM.

Methyl ester derivatives of carboxylic acid possess a higher BBB permeability (Shukuri et al., 2011; Suzuki et al., 2004; Takashima-Hirano et al., 2010). Thus, we investigated whether phenylacetic acid methyl ester (PhAcM) works as a ligand precursor that is efficiently delivered into the brain through the BBB and converted to PhAc by an endogenous esterase activity, resulting in the activation of target neurons (see Figure 1A for a model of drug delivery). We administered PhAcM solution (20 mg/kg) or vehicle intravenously into a lateral tail vein of the Tg mice and monitored the firing rate of the LC neurons using an extracellular recording. NE cells in the LC were identified as described previously (Fukabori et al., 2020). The normalized firing rate was significantly elevated following PhAcM administration (Figure 1B; post vs. pre, one-way ANOVA, F_(1, 10)_ = 5.62, p = 0.039, partial η^2^ = 0.36), but not following vehicle administration (F_(1, 11)_ = 2.63, p = 0.133, partial η^2^ = 0.19).

Next, we examined changes in NE_[ext]_ in the ACC of the Tg mice by the peripheral PhAcM administration. PhAcM administration resulted in a marked increase in NE_[ext]_ in the Tg mice (Figure 1C; two-way ANOVA, group effect: F_(2, 9)_ = 24.95, p < 0.001, partial η^2^ = 0.84, fraction effect: F_(6, 54)_ = 6.81, p < 0.001, partial η^2^ = 0.43, interaction: F_(12, 54)_ = 5.58, p < 0.001, partial η^2^ = 0.55) These data show that peripheral administration of methyl ester of PhAc efficiently stimulates the IR84a/IR8a-expressing target neurons in the brain.

### Processing of the ligand precursor results in the activation of the IR84a/IR8a-expressing neurons in the brain

Our *in vivo* electrophysiological and biochemical experiments demonstrate that peripherally administered PhAcM induces excitation of the central IR84a/IR8a-expressing neurons (Figure 1B and C), suggesting that a precursor of the ligand is converted to the ligand by endogenous esterase following translocation across the BBB. To test this possibility, we synthesized a ^11^C-incorporated analogue of PhAcM, 2-phenyl[3-^11^C]propionic acid methyl ester ([^11^C]PhPrM) in which the methyl group at the 3-position of 2-phenylpropionic acid methyl ester (PhPrM) was labeled with ^11^C. Although there were two stereoisomers with the (*S*)– and (*R*)-configurations regarding 2-phenylpropionic acid (PhPr), we demonstrated that only (*S*)-PhPr can activate the IR84a/IR8a-expressing neurons by an *ex vivo* whole-cell current clamp recording and *in vivo* microdialysis with the LC-NE system of the Tg mice (Figure 2—figure supplement 1). Thus, we modeled that peripherally administered the methyl ester form (*S*)-PhPrM crosses the BBB and enters the brain, where (*S*)-PhPrM is metabolized to its acid form (*S*)-PhPr by esterase activity (Figure 2A). We injected (*S*)-[^11^C]PhPrM into a lateral tail vein of the Tg mice and removed their brains 30 min later. Tissue suspension was prepared from the LC region and subjected to thin-layer chromatography (TLC) analysis (Figure 2—figure supplement 2). The results showed the presence of (*S*)-[^11^C]PhPr in the LC tissue, supporting the model for the translocation of peripherally administered (*S*)-[^11^C]PhPrM across the BBB into the brain and the following conversion to (*S*)-[^11^C]PhPr by esterase activity.

**Figure 2.**
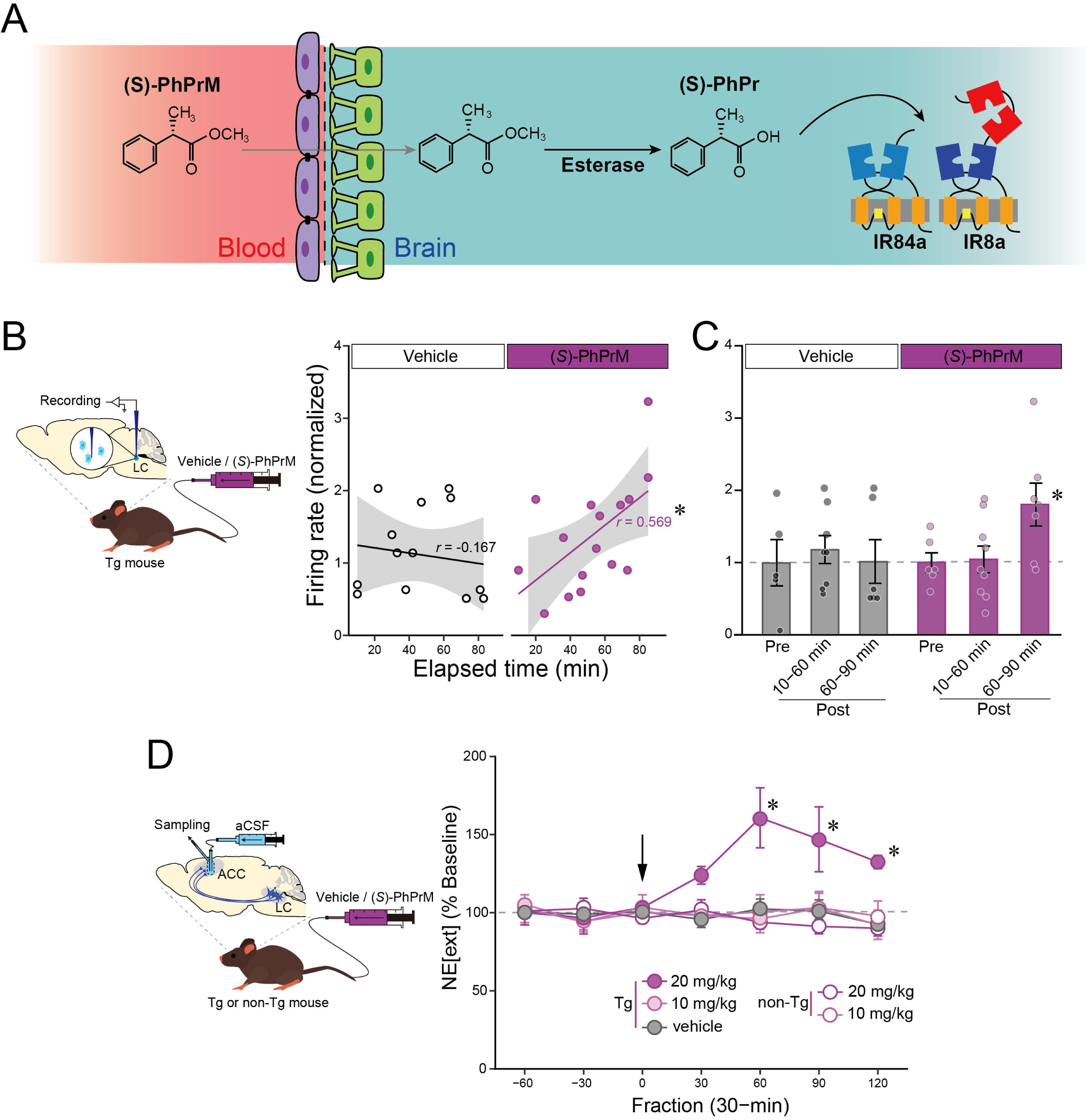
Identification of (*S*)-PhPr as an IR84a/IR8a ligand and enhancement of the BBB permeability of (*S*)-PhPr by methyl esterification. (**A**) Schematic representation of enhanced BBB permeability of PhPr. It was expected that peripherally administered methyl ester of PhPr (PhPrM) penetrates the brain across the BBB and converted PhPr stimulates the IR84a/IR8a complex. (**B**) Plot of correlation between the normalized firing rate of the LC neurons and the elapsed time (min) after (*S*)-PhPrM (20 mg/kg) or vehicle tail vein injection in the Tg mice (vehicle n = 14, (*S*)-PhPrM n = 16). *p < 0.05. (**C**) Plot of changes in the normalized firing rate by the (*S*)-PhPrM (20 mg/kg) and vehicle tail vein injection in the Tg mice (vehicle/pre n = 5, vehicle/post 10-60 min n = 8, vehicle/post 60-90 min n = 6, (*S*)-PhPrM/pre n = 6, (*S*)-PhPrM/post 10-60 min n = 9, (*S*)-PhPrM/post 60-90 min n = 7). *p < 0.05 vs. (*S*)-PhPrM/pre. Data are presented as mean ± SEM and individual data points are overlaid. (**D**) Plot of changes in the extracellular NE level (NE_[ext]_) in the ACC of the Tg and non-Tg mice before and after (*S*)-PhPrM (10 or 20 mg/kg) tail vein injection (n = 3 for all groups). NE_[ext]_ is expressed as a percentage of the average baseline levels of each mouse. Arrow indicates the timing of drug injection. At the 60– to 120-min fractions, NE_[ext]_ of 20 mg/kg (*S*)-PhPrM-administered Tg mice was significantly higher than that of vehicle-(ts_(70)_ = 5.18, 4.08, 3.55, ps < 0.001, rs = 0.51, 0.44, 0.39, respectively) or 10 mg/kg (*S*)-PhPrM-administered Tg mice (ts_(70)_ = 5.75, 3.93, 3.57, ps < 0.001, p = 0.001, rs = 0.57, 0.43, 0.39, respectively) and that of 10 mg/kg (ts_(70)_ = 5.36, 3.90, 3.07, ps < 0.001, p = 0.003, rs = 0.54, 0.42, 0.34, respectively) or 20 mg/kg (*S*)-PhPrM-administered non-Tg littermates (ts_(70)_ = 5.94, 4.96, 3.77, ps < 0.001, rs = 0.58, 0.51, 0.41, respectively).*p < 0.05 vs. other four groups. Data are presented as mean ± SEM.

We next tested whether peripherally administered (*S*)-PhPrM stimulates central noradrenergic activity in the Tg mice using *in vivo* extracellular recording. The normalized firing rate of the LC-NE neurons showed a gradual increase following the injection of (*S*)-PhPrM (20 mg/kg) into the tail vein of the Tg mice, whereas no such increase was observed following vehicle injection (Figure 2B); statistical test supported the observation, and Pearson correlation (r) between the elapsed time after drug administration and firing rate (normalized) was significant for (*S*)-PhPrM (r = 0.569, p = 0.021) but not for vehicle (r = –0.167, p = 0.567). When we divided the data into three time periods (pre-administration, 10-60 min after the administration, and 60-90 min after the administration) and compared the mean firing rate for each period (Figure 2C), the mean firing rate up to 60 min after (*S*)-PhPrM administration (10-60 min) was not significantly different from that of pre-administration (one-way ANOVA, F_(1, 13)_ = 0.03, p = 0.872, partial η^2^ = 0.002), but the mean firing rate after 60 min of (*S*)-PhPrM administration (60-90 min) was significantly higher than that of pre-administration (F_(1, 11)_ = 5.38, p = 0.041, partial η^2^ = 0.33). In the vehicle condition, there were no significant differences between pre vs. 10-60-min post-administration (F_(1, 11)_ = 0.27, p = 0.614, partial η^2^ = 0.02) and pre vs. 60-90-min post-administration (F_(1, 9)_ = 0.001, p = 0.970, partial η^2^ = 0.0001).

Furthermore, we examined the change in NE_[ext]_ in the ACC of the Tg and non-Tg mice by the peripheral (*S*)-PhPrM administration (Figure 2D). A 20 mg/kg (*S*)-PhPrM resulted in a slightly delayed but long-lasting increase in NE_[ext]_ in the Tg mice (two-way ANOVA, group effect: F_(4, 10)_ = 9.73, p = 0.002, partial η^2^ = 0.80, fraction effect: F_(6, 24)_ = 1.93, p = 0.090, partial η^2^ = 0.16, interaction: F_(6, 60)_ = 2.45, p = 0.003, partial η^2^ = 0.49) These data show that the peripherally administered ligand precursor, (*S*)-PhPrM, is converted to (*S*)-PhPr in the brain following translocation across the BBB, which stimulates the IR84a/IR8a-expressing LC-NE neurons.

### Functional expression of IR84a/IR8a in specific cell types by using a viral vector system

We examined whether the IRNA technology could be applied to non-NE neurons in species other than mice using a viral vector strategy carrying the flip-excision switch (FLEX) system (Kato et al., 2018; Kato et al., 2007; Matsushita et al., 2023). For this aim, we employed the striatum of rats as a model. The striatum controls behaviors through the activity distributed across two subpopulations of GABAergic spiny projection neurons (SPNs), which express distinct DA receptor subtypes that respond to DA in an opposing manner (Alexander and Crutcher, 1990; Gerfen and Bolam, 2016): direct SPNs (dSPNs) express type 1 receptors (D1Rs) and project to the substantia nigra pars reticulata and entopedoncular nucleus; indirect SPN (iSPNs) express type 2 receptors (D2Rs), together with adenosine A_2A_ receptor (A_2A_R) (Fink et al., 1992; Schiffmann et al., 1991), sending axons to the external segment of the globus pallidus (GPe). We employed *Drd2*-Cre transgenic rats that express Cre recombinase predominantly in iSPNs (Nonomura et al., 2018). Triple immunohistochemistry of the section through the striatum of the *Drd2*-Cre rats for Cre, A_2A_R, and D1R established that the Cre transgene was highly and specifically expressed in iSPNs but was absent in dSPNs (Figure 3—figure supplement 1). A lentiviral vector pseudotyped with vesicular stomatitis virus glycoprotein (VSV-G) was constructed to express EGFP/IR84a-2A-IR8a in a Cre-dependent manner (Lenti-FLEX-EGFP/IR84a-2A-IR8a) and then microinjected into the dorsal striatum of the *Drd2*-Cre rats (Figure 3A). Striatal sections were stained by triple immunohistochemistry for GFP, IR8a, and A_2A_R, or GFP, IR8a, and D1R (Figure 3A). The numbers of immuno-positive cells (total GFP^+^, total IR8a^+^, GFP^+^ & IR8a^+^, total A_2A_R^+^, total D1R^+^, GFP^+^ & A_2A_R^+^, GFP^+^ & D1R+, IR8a^+^ & A_2A_R^+^, IR8a^+^ & D1R^+^, GFP^+^ & IR8a^+^ & A_2A_R^+^, GFP^+^ & IR8a^+^ & D1R^+^) were counted (Supplementary Table 1). The frequencies of EGFP/IR84a and IR8a double-expression in two types of neurons (efficiency) were 73.92 ± 0.01% in A_2A_R-positive neurons and 2.52 ± 0.01% in D1R-positive neurons (n = 4 animals), frequencies of two types of neurons in EGFP/IR84a and IR8a double-expressing neurons (specificity) were 79.87 ± 0.03% for A_2A_R-positive neurons and 2.28 ± 0.01% for D1R-positive neurons (n = 4 animals), indicating the efficient and specific expression of EGFP/IR84a and IR8a in striatal iSPNs in the viral vector-injected *Drd2*-Cre rats.

**Figure 3.**
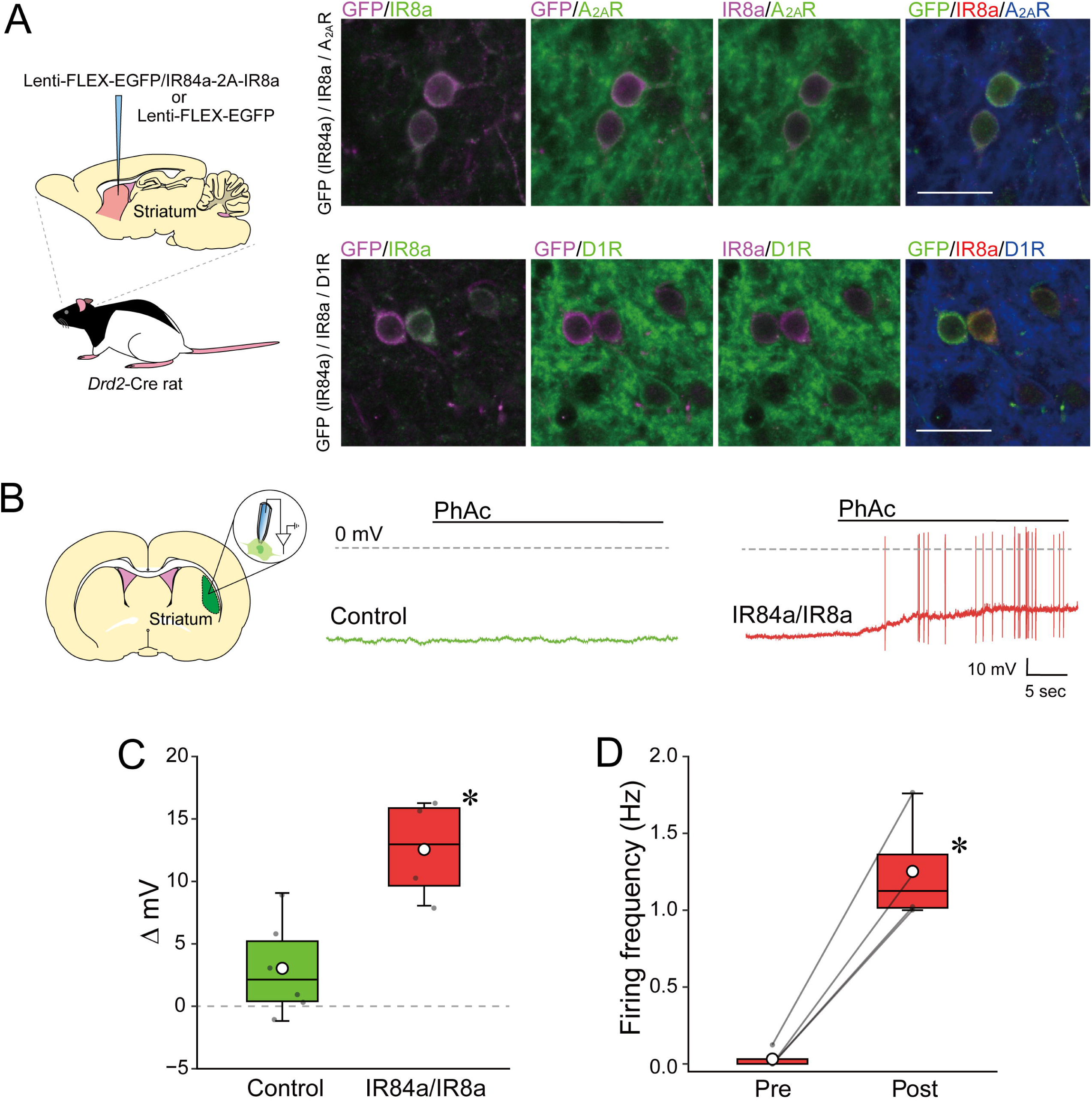
Cell type-specific expression of IR84a/IR8a in the rat striatum and responses to PhAc. (**A**) Robust expressions of IR84a and IR8a in iSPNs (A_2A_R+), but not in dSPNs (D1R+) in Tg rats. A lentiviral vector expressing EGFP/IR84a and IR8a in a Cre-dependent manner was microinjected into the striatum of the *Drd2*-Cre rats. Sections were triple-stained through immunohistochemistry for GFP, IR8a, and A_2A_R, or GFP, IR8a, and D1R. Scale bar: 25 um. (**B**) Representative whole-cell recordings of the Lenti-FLEX-EGFP (control) or Lenti-FLEX-EGFP/IR84a-2A-IR8a (IR84a/IR8a) vector-treated striatal neurons of the *Drd2*-Cre rats in response to PhAc. (**C-D**) Plot of changes in the membrane potential (**C**) and firing frequency (**D**) with addition of PhAc (0.1% w/v) in the control (n = 6) and IR84a/IR8a-expressing neurons (n = 4). Individual data points are overlaid *p < 0.05. Each white dot in the boxplot indicates the mean for the condition.

To determine whether striatal cells expressing IR84a/IR8a show neuronal activation by PhAc, we induced expression of either both EGFP/IR84a and IR8a or GFP (as the control condition) in striatal iSPNs using the lentiviral vector system and performed a whole-cell current clamp recording in slice preparations (Figure 3B). The amount of difference in the membrane potential between pre– and post-PhAc (0.1%) bath application (Δ mV) of striatal neurons expressing IR84a and IR8a was significantly greater than that of the control neurons expressing GFP but not IR84a/IR8a (Figure 3C; one-way ANOVA, genotype effect: F_(1, 8)_ = 14.03, p = 0.006, partial η^2^ = 0.64). The IR84a/IR8a expressing neurons also showed significant elevation in the firing rate (normalized) by PhAc application (Figure 3D; one-way ANOVA, genotype effect: F_(1, 3)_ = 69.61, p = 0.004, partial η^2^ = 0.96). These data suggest that the viral vector gene expression system allows a cell type-specific, highly efficient expression of IR84a/IR8a and that cells expressing the two subunits show neuronal activation in a ligand-dependent manner.

### Peripheral administration of the ligand precursors induces excitation of target neurons expressing IR84/IR8a with a viral vector system

Striatal neurons of rats expressing IR84a/IR8a by the viral vector system were responsive to PhAc in the slice preparation. We then investigated whether these neurons respond to peripherally administered ligand precursors *in vivo*. We expressed EGFP/IR84a and IR8a in the iSPNs in the unilateral striatum (left or right) of the *Drd2-Cre* rats by microinjection of the lentiviral vector and then examined whether a cocaine-induced DAergic activation of the striatum is modulated by the peripherally administered PhAcM, resulting in a rotation behavior to a specific direction (Figure 4A). The rats showed contraversive rotation when PhAcM (10 mg/kg) was administered intraperitoneally (i.p.) but did not show such a directional bias when the vehicle was administered (Figure 4B; three-way ANOVA, drug effect: F_(1, 7)_ = 0.45, p = 0.542, partial η^2^ = 0.06, direction effect: F_(1, 7)_ = 0.09, p = 0.780, partial η^2^ = 0.01, time-bin effect: F_(11, 77)_ = 1.48, p = 0.156, partial η^2^ = 0.17, drug × direction interaction: F_(1, 7)_ = 9.49, p = 0.018, partial η^2^ = 0.58, drug × time-bin interaction: F_(11, 77)_ = 0.33, p = 0.977, partial η^2^ = 0.04, direction × time-bin interaction: F_(11, 77)_ = 0.82, p = 0.620, partial η^2^ = 0.10, drug × direction × time-bin interaction: F_(11, 77)_ = 2.22, p = 0.022, partial η^2^ = 0.24); the number of contraversive rotation in the PhAcM administration was significantly greater than that in the vehicle administration in the first and second 5-min bin (Fs_(1, 168)_ = 5.04, 5.44, ps = 0.026, 0.021, partial η^2^s = 0.21, 0.23).

**Figure 4.**
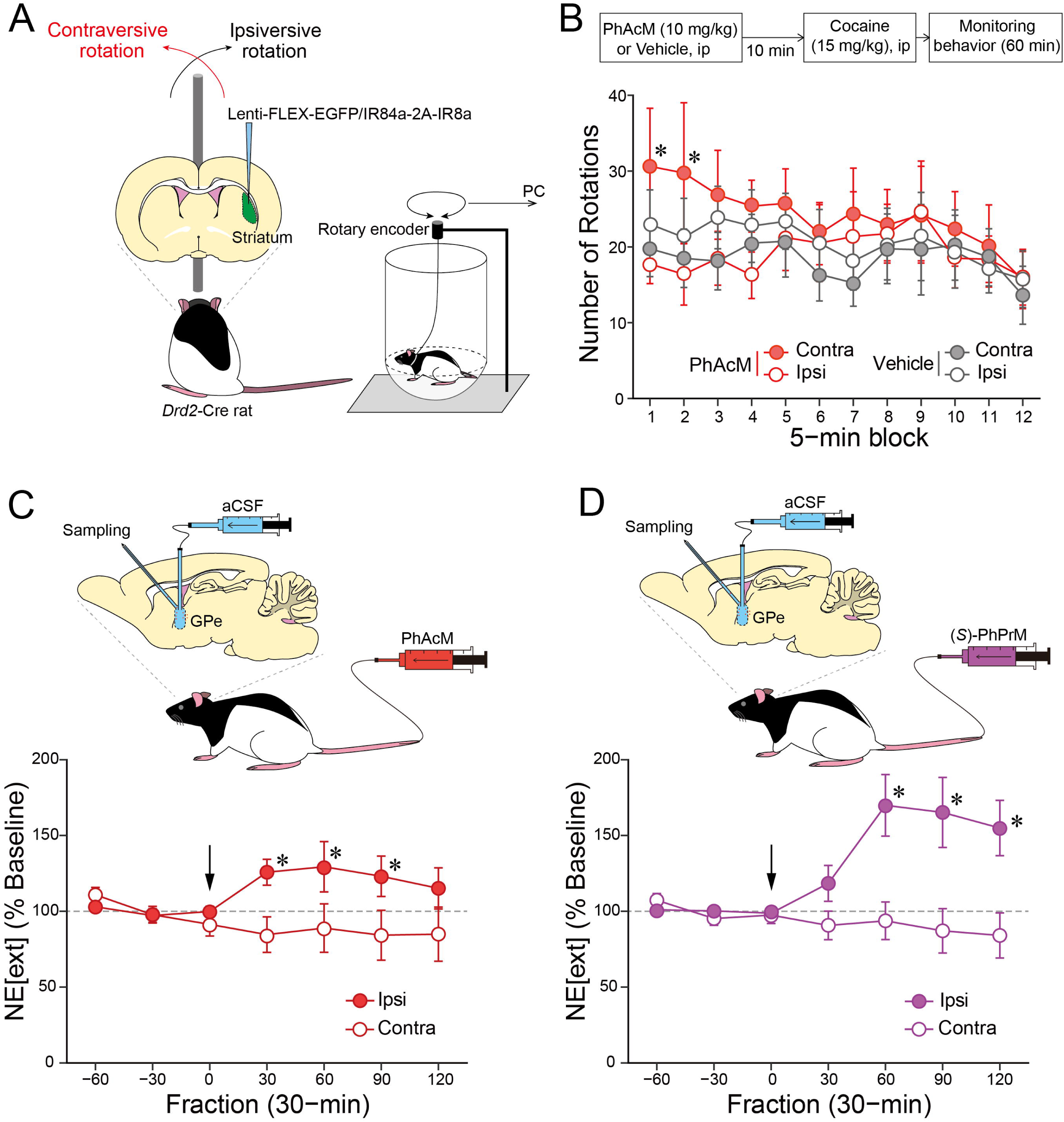
Effects of peripherally administered ligand-dependent activation of the IR84a/IR8a expressing iSPNs. (**A**) Schematic representation of behavioral paradigm to monitor PhAcM-dependent motor effects. The *Drd2*-Cre rats received the lentiviral vector that induces IR84a/IR8a in a Cre-dependent manner in the unilateral striatum. Their rotational behavior was monitored in a cylinder with hemispheric bottom via a rotary encoder connected to PC. (**B**) Schedule of the rotational behavior test (upper panel). Rats were ip injected with PhAcM (10 mg/kg) or vehicle, and then cocaine after a 10-min interval, followed by behavioral monitoring in the apparatus. Plot of changes in the rats’ ipsiversive and contraversive rotations induced by cocaine following PhAcM injection (lower panel) (ns = 8 for vehicle and PhAcM conditions). *p < 0.05 vs. PhAcM/ipsi. (**C**) Plot of changes in the extracellular GABA level (GABA_[ext]_) in the GPe of the viral vector-treated *Drd2*-Cre rats before and after PhAcM (10 mg/kg) tail vein injection. Ipsilateral and contralateral GPes of the viral vector-treated striatum were compared (n = 5). GABA_[ext]_ in the ipsilateral GPe was significantly higher than that of the contralateral GPe at the 30– to 90-min fractions (Fs_(1, 56)_ = 6.42, 6.19, 5.73, ps = 0.014, 0.016, 0.020, partial η^2^ = 0.014, 0.016, 0.020, respectively). *p < 0.05. (**D**) Plot of changes in GABA_[ext]_ in the GPe of the viral vector-treated *Drd2*-Cre rats before and after PhPrM (20 mg/kg) tail vein injection. Ipsilateral and contralateral GPes of the viral vector-treated striatum were compared (n = 4). GABA_[ext]_ in the ipsilateral GPe was significantly higher than that in the contralateral GPe at the 60– to 120-min fractions (Fs_(1, 42)_ = 13.32, 17.34, 11.19, p = 0.001, p < 0.001, p = 0.002, partial η^2^ = 0.37, 0.43, 0.33, respectively). *p < 0.05. GABA_[ext]_ is expressed as a percentage of the average baseline levels of each rats. Arrows indicate the timing of drug injection. Data are presented as mean ± SEM.

We then examined the change in GABA release in the GPe by the intravenous administration of (*S*)-PhPrM (20 mg/kg) as well as PhAcM (10 mg/kg) with the rats completing the behavioral experiment. The effects of these drugs on extracellular GABA level (GABA_[ext]_) were assessed in the ipsilateral GPe to the viral vector-injected striatum with the contralateral GPe (ipsilateral to the intact striatum) as the control condition. PhAcM administration resulted in a rapid and sustained increase in GABA_[ext]_ in the ipsilateral GPe relative to the contralateral GPe (Figure 4C; two-way ANOVA, hemisphere effect: F_(1, 8)_ = 3.76, p = 0.089, partial η^2^ = 0.32, fraction effect: F_(6, 48)_ = 0.61, p = 0.719, partial η^2^ = 0.07, interaction: F_(6, 48)_ = 2.69, p = 0.025, partial η^2^ = 0.25). (*S*)-PhPrM resulted in a slightly delayed but long-lasting increase in GABA_[ext]_ in the ipsilateral GPe relative to the contralateral GPe (Figure 4D; two-way ANOVA, hemisphere effect: F_(1, 6)_ = 7.46, p = 0.034, partial η^2^ = 0.55, fraction effect: F_(6, 36)_ = 1.76, p = 0.135, partial η^2^ = 0.23, interaction: F_(6, 36)_ = 5.66, p < 0.001, partial η^2^ = 0.49). These behavioral and biochemical data show that the target neurons expressing IR84a/IR8a based on the viral vector gene expression system respond to peripherally administered ligand precursors *in vivo*.

### Visualization of radiolabeled ligand binding on IR84a/IR8a-expressing neurons following peripheral administration of the methyl ester precursor

Striatal iSPNs expressing IR84a/IR8a using the viral vector system revealed the responsiveness to the peripheral administration with (*S*)-PhPrM and PhAcM. Finally, we sought to visualize the binding of (*S*)-PhPr, converted in the brain from peripherally administered (*S*)-PhPrM, to the IR84a/IR8a complex in striatal neurons. (*S*)-[^11^C]PhPrM was administered intravenously into the tail vein of the *Drd2*-Cre rats that had been microinjected unilaterally with a cocktail of AAV2-EF1α-FLEX-EGFP/IR84a and AAV2-EF1α-FLEX-HA/IR8a vectors into the striatum, and to image ^11^C in the striatum *in vivo* using positron emission tomography (PET) and *ex vivo* using autoradiography (ARG) (Figure 5A). PET imaging showed a high level of ^11^C radioactivity in the viral vector-injected side of striatal regions of (*S*)-[^11^C]PhPrM-administered rats (Figure 5B). Similarly, *ex vivo* ARG showed a high accumulation of ^11^C radioactivity in the vector-treated side of the striatum of the same animals (Figure 5C). Based on the ARG data, we compared ^11^C radioactivity [(PSL/mm^2^)/(MBq/kg)] between the vector-treated and intact sides of the striatum, and the vector-treated side had greater ^11^C radioactivity than the intact side (Figure 5D; one-way ANOVA, the main effect of side: F_(1, 16)_ = 44.23, p < 0.001, partial η^2^ = 0.73). Since metabolite analyses by TLC (Figure 2—figure supplement 2) showed that most of the (*S*)-[^11^C]PhPrM was converted to (*S*)-[^11^C]PhPr in the brain within at least 30 min after the administration, the ^11^C radioactivity visualized in the PET and ARG was considered to be derived from (*S*)-[^11^C]PhPr, but not from (*S*)-[^11^C]PhPrM. Thus, these data support the binding of the ligand that was converted from the systemically administered precursor to the IR84a/IR8a complex in striatal neurons.

**Figure 5.**
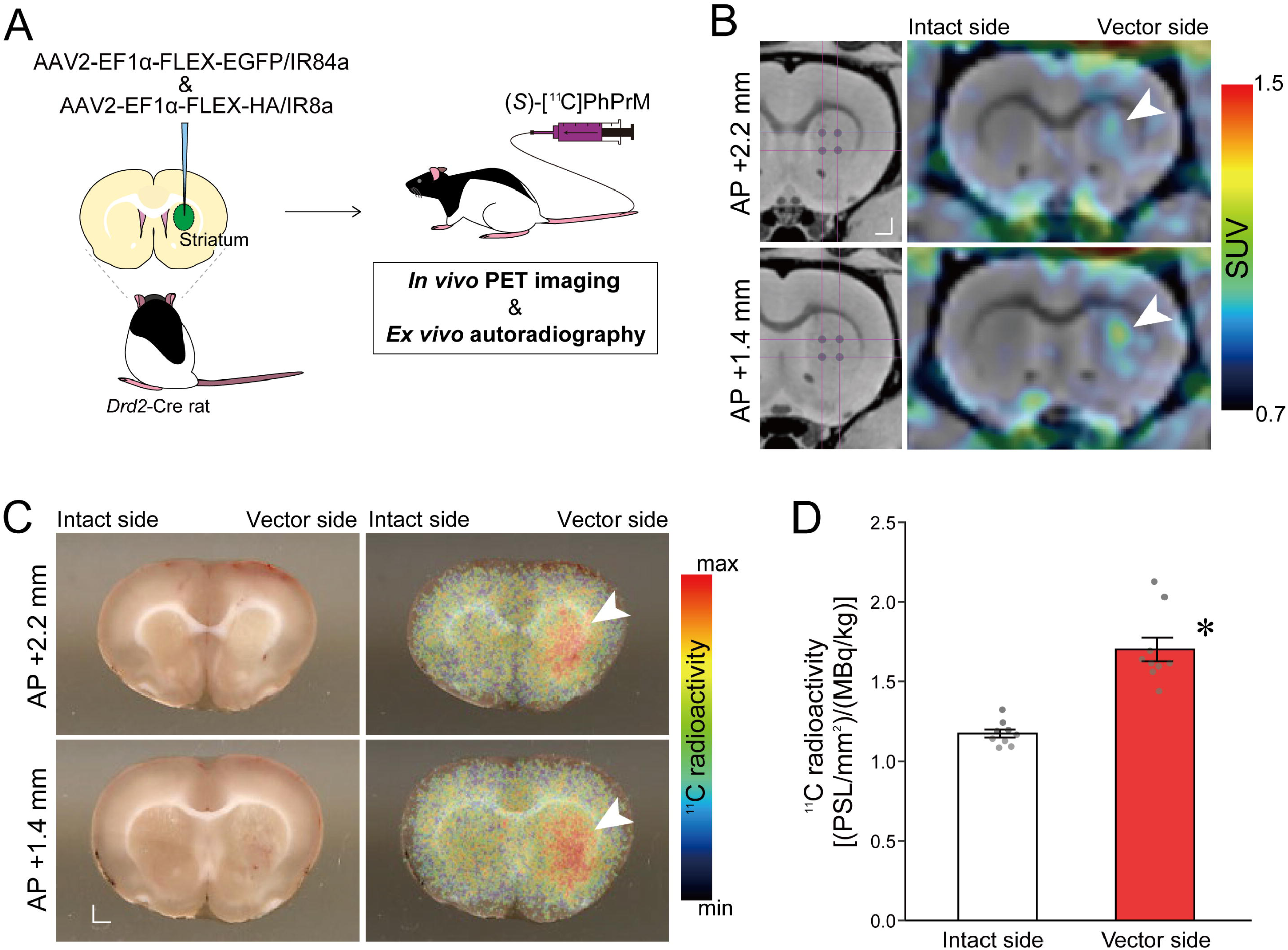
Visualization of the bindings of (*S*)-PhPr on the IR84a/IR8a expressing neurons following peripheral administration of the methyl ester precursor. (**A**) Schematic representation of brain imaging. The *Drd2*-Cre rats received the unilateral injection of AAV2 vectors that induce IR84a and IR8a in a Cre-dependent manner. These rats were administered (*S*)-[^11^C]PhPrM into the tail vein and used for in *vivo* PET imaging and *ex vivo* ARG.(**B**) Representative coronal PET images with ^11^C overlaid on the magnetic resonance T1-weighted images of the viral vector-treated *Drd2*-Cre rat. Corresponding sites of viral vector injections were indicated in the left panel. (**C**) Representative ARG images with ^11^C for the coronal sections of the vector-treated *Drd2*-Cre rat. Corresponding bright-field images of the sections were indicated in the left panel. The arrowheads indicate the high level ^11^C radioactivity in the striatal regions. Scale bar: 1 mm (**B-C**). (**D**) Plot of ^11^C radioactivity quantified from the ARG data compared between the viral vector-treated and intact sides of the striatum (ns = 9). *p < 0.05. Data are presented as mean ± SEM and individual data points are overlaid.

## Discussion

This study aimed to improve the previously reported mammalian neural activation technology employing insect IRs (Fukabori et al., 2020) in three ways: first, through development of a pro-drug system that is administered peripherally and then transferred to the central nervous system to induce activation in the target cells expressing IR84a and IR8a, second, a viral vector system that induces expression of the IR84a/IR8a complex in target cell populations, and third, to generalize the functional application of IRNA to other neuron types and mammalian species. Since it was difficult to deliver PhAc into the brain across the BBB, intracranial microinjection of the ligand was necessary to activate the IR84a/IR8a expressing cells in the brain (Fukabori et al., 2020). We therefore developed a strategy to activate the target neurons by esterifying PhAc to make it lipophilic, which facilitates the delivery through the BBB by passive diffusion and reverts it to the activated form by esterase activities in the brain. Moreover, we generated viral vectors that induce IR84a/IR8a expression according to the Cre-loxP system and microinjected them into the striatum of the *Drd2*-Cre rats. We found striatal iSPNs-specific IR84a/IR8a expression, an excitatory response of iSPNs to PhAc *ex vivo*, behavioral and biochemical responses of iSPNs to PhAcM *in vivo*, and accumulation of labeled ligands at the vector injection site, demonstrating the utility of the viral vector system in expressing functional IR84a/IR8a complex to Cre-expressing cells of Tg animals.

In comparison with the peripheral administration of PhAcM (Figure 1B and C), the peripherally administered (*S*)-PhPrM revealed a delayed noradrenergic activation in the Tg mice, both in the electrophysiological activity of the LC neurons (Figure 2B and C) and the NE release in the terminal region (Figure 2D). Similar delayed response of the IR84a/IR8a-expressing neurons to (*S*)-PhPrM was observed in the striatal neurons of rats, in which the GABA release in the terminal region of the IR84a/IR8a expressing iSPNs that was rapidly elevated following PhAcM administration rose slowly following (*S*)-PhPrM (Figure 4C and D). It should be noted that (*S*)-PhPr induced a rapid, excitatory responses in IR84a/IR8a-expressing neurons in the acute LC slice of the Tg mice (Figure 2—figure supplement 1A-C). The delayed effects of (*S*)-PhPrM may be attributable to the reduced efficiency of the hydrolysis by conformational interference in the active site of esterase (Imai et al., 2006), resulting in the change in time course of (*S*)-PhPr bioavailability in the target brain regions.

In our previous report, we used the IRNA technique to clarify the role of LC-NE neurons in emotional memory processing (Fukabori et al., 2020). The present study extends the utility of IRNA as a tool for elucidating the neural basis of behavior by analyzing the role of striatal iSPNs in movement control. When specific activation of iSPNs was induced by IRNA system in the unilateral striatum, contraversive rotations were apparent after systemic administration of cocaine. A report that systemic amphetamine administration after selective ablation of iSPNs in the unilateral striatum produced ipsilateral rotation of mice (Sano et al., 2003) is in line with the present results. With respect to spontaneous movement, unilateral activations of striatal iSPNs by optogenetic (Grimm et al., 2021; Kravitz et al., 2010; Lee et al., 2016) or DREADD-based chemogenetic (Alcacer et al., 2017) were reported to induce the ipsiversive rotations in mice. Therefore, the function of iSPNs in the movement control seems to change between spontaneous and drug-induced conditions. Besides, several studies investigating the effects of unilateral striatal excitotoxic lesion and subsequent psychostimulant administration in rats reported rotations in both directions or no rotations (Dunnett & Iversen, 1982; Dunnett et al., 1988; Norman et al., 1988). The reliability and direction of rotations were dependent on the topography of the lesion sites within the striatum (Fricker et al., 1996; Norman et al., 1992). It should be noted that the present study induced activation in iSPNs in the relatively caudolateral region of the striatum (Figure 4—figure supplement 1A). Systematic studies on the effects of remote manipulation of iSPN activities in various striatal subregions on drug-induced and spontaneous behaviors are needed in the future.

The IRNA technology is based on insect-derived environmental sensing receptors (Croset et al., 2010; Depetris-Chauvin et al., 2015), a highly divergent subfamily of iGluR, which has not been developed as chemogenetic tools. The invertebrate origin of IRNA’s genetic components allows us to omit the process of producing knockout background animals (Atasoy and Sternson, 2018; Lima and Miesenböck, 2005), as is the case of the chemogenetic systems based on P2X (Zemelman et al., 2003) and TRP channels (Arenkiel et al., 2008; Güler et al., 2012). Furthermore, compared to G protein-coupled receptor-based chemogenetic systems that depend on the intrinsic second messenger pathways within the targeted cells, the operating mechanism of IRNA is considered far more uncomplicated and is expected to open the way for remote manipulation of targeting cells that have been difficult to approach with existing technologies (Fukabori et al., 2020; Prieto-Godino et al., 2021).

Since molecules that can be sensed varies dramatically by changing the subunit composition of the IR complex (Abuin et al., 2011; Ni, 2020; van Giesen and Garrity, 2017; Wicher and Miazzi, 2021), the use of new receptor complexes other than the IR84a/IR8a is feasible for the application of IRNA technology. In addition, searching for actuator molecules for IR84a/IR8a with higher affinity than PhAc, PhAl, and (*S*)-PhPr would also lead to a more efficient chemogenetic system. Nevertheless, having shown that IRNA works in several cell types in two different mammalian systems, our study indicate the potential of this system in future application as a chemogenetic-based therapeutic strategy for neurological and neuropsychiatric diseases (Urban and Roth, 2015; Walker and Kullmann, 2020).

## Materials and methods

### Animals

Tg mice including B6.Cg-Tg(TH-GFP-IR84a/IR8a)2-1Koba (Fukabori et al., 2020) and B6.B6D2-Tg(Th-EGFP)21-31Koba (Matsushita et al., 2002) (Riken BioResource Research Center) were bred with wild-type C57BL/6J mice (CLEA Japan) to produce the heterozygous and wild-type offspring. LE-Tg(Drd2-cre)490-9Koba Tg rats (Nonomura et al., 2018) (The National BioResource Project for the Rat in Japan) were mated with wild-type Long Evans rats (Charles River Laboratories) to obtain the heterozygous and wild-type offspring. Animal care and handling procedures followed the guidelines established by the Laboratory Animal Research Center of Fukushima Medical University and RIKEN Biosystems Dynamics Research. All procedures were approved by the Fukushima Medical University Institutional Animal Care and Use Committee and RIKEN Biosystems Dynamics Research. Mice and rats were maintained on a 12 h light/dark cycle (lights on at 07:00 h) at an ambient temperature of 22°C. All experimental procedures were conducted during the light period. Mice aged 12–14 weeks old and rats aged 8–16 weeks old were used for the following experiments. Male rats were used in the behavioral and biochemical experiment. In all other experiments, both males and females were used. Mice were housed in groups of three to five, whereas rats were housed in groups of two to three. They were single-housed after the stereotaxic surgeries for the microdialysis experiments. The assignment of animals to the experimental conditions was random.

### Chemical compounds

Phenylacetic acid (PhAc, 166-01232; Wako Pure Chemical) was dissolved in PBS and used as the ligand of the IR84a/IR8a complex to induce neuronal activation. For peripheral administration, Phenylacetic acid methyl ester (PhAcM, 108057; Sigma-Aldrich) and (*S*)-2-phenylpropionic acid methyl ester ((*S*)-PhPrM; Sumika Technoservice) were dissolved in PBS containing 1.5% Tween80 and 2.5% ethanol (vehicle) and administered in the lateral tail vein of mice and rats (at 5 ml/kg) or i.p. injected in rats (at 5 ml/kg). For intracranial microinjection for mice, (*S*)-2-phenylpropionic acid ((*S*)-PhPr, 279900; Sigma-Aldrich), (*R*)-2-phenylpropionic acid ((*R*)-PhPr, 279897; Sigma-Aldrich) were dissolved in the PBS at a concentration of 0.6% (w/v). For the synthesis of (*S*)-2-phenyl[3-^11^C]propionic acid methyl ester ((*S*)-[^11^C]PhPrM), our previous radiolabeling method was used (Takashima-Hirano, et al., 2010). (*R*),(*S*)-[^11^C]PhPrM was synthesized from the precursor phenyl-propionate by the reaction of [^11^C]CH_3_I with 1 M tetrabutylammonium fluoride in the presence of anhydrous dimethyl sulfoxide, and subsequent separation of racemic form to (*S*)-[^11^C]PhPrM by a chiral column (CHIRALPAK OJ-RH, DAICEL, Japan). The radioactivity at the time of the radiosynthesis was 1,990-3,160 MBq (molar radioactivity 28-73 GBq/μmol), and the chemical and radiochemical purities were over 99%. In the behavioral experiments, cocaine hydrochloride (Takeda Pharmaceutical) dissolved in saline was i.p. administrated at 15 mg/kg in rats.

### TLC analysis

The brain was removed from the Tg mice under deep anesthesia combined with pentobarbital and isoflurane 30 minutes after intravenous administration of the (*S*)-[^11^C]PhPrM (42 MBq/0.1 ml). The brain tissue was homogenized with a five-fold volume of physiological saline on ice and then vortexed with an equal volume of acetonitrile. The samples were centrifuged (9700 × *g*, for 2 min, at 4°C), and the supernatant obtained was spotted 2 μl on the origin point of a reversed-phase plate (RP-18, Merck Millipore) and developed with a mixture solution of acetonitrile, distilled water, and formic acid (65:34:1). To further determine whether the metabolite detected by radio-TLC was (*S*)-PhPr which is the active form to bind IR receptors, we applied with unlabeled (*S*)-PhPrM and (*S*)-PhPr solutions to the origin point on a reversed-phase plate coated with fluorescent indicator (RP-18 F254S, Merck Millipore). The developed spots were detected with ultraviolet light (254 nm).

### Viral vector preparation

Lentiviral vectors pseudotyped with vesicular stomatitis virus glycoprotein (VSV-G) were prepared as previously described (Kato et al., 2007) with some modifications. The GFP-tagged IR84a and IR8a genes were connected by 2A peptide sequence to construct the flip-excision switch (FLEX) system (FLEX-EGFP/IR84a-2A-IR8a). Then, a transfer plasmid containing the FLEX-EGFP/IR84a-2A-IR8a downstream of the mouse stem cell virus promoter and an envelope plasmid under the control of the cytomegalovirus enhancer/chicken β-actin promoter, VSV-G cDNA. HEK293T cells were transfected with transfer, envelope, and packaging plasmids using calcium phosphate precipitation. Viral vector particles were pelleted by centrifugation at 6,000 × *g* for 16-18 h and suspended in phosphate-buffered saline (PBS); different concentrations (55% and 20%) of sucrose in PBS and particle solution were successively layered from the bottom of the ultracentrifuge tube. The tubes were centrifuged at 100,000 × *g* for 2 hours using a SW 55Ti swinging-bucket rotor (Beckman Coulter).

Adeno-associated virus (AAV) vector was prepared using the AAV Helper-Free system (Agilent Technologies) as described previously (Kato et al., 2018; Matsushita et al., 2023). HEK293T cells (American Type Culture Collection) were transfected with plasmids encoding transfer genes (FLEX-IR84a/EGFP and FLEX-hemagglutinin (HA)/IR8a) downstream of the elongation factor 1α promoter, adeno-helper genes, and adenovirus genes required for AAV replication and encapsulation by a calcium phosphate precipitation method. After 48 hours of transfection, cells were collected, lysed, and crude viral vector lysate was purified with the first round of CsCl gradient ultracentrifugation by using a NVT 65 near-vertical tube rotor (Beckman Coulter) with a quick-seal tube at 287,000 × *g* for 24 h at 16°C. The tube was inserted by a needle into the bottom to collect drop solution of 1.0 mL into a tube (1-10 fractions). To confirm the peak fractions, real-time quantitative PCR was performed using each fraction as a template. The second round of CsCl gradient ultracentrifugation was performed using an SW 55Ti rotor (Beckman Coulter) with a thinwall polypropylene tube at 287,000 × *g* for 24 h at 16°C. Drop solution of 0.5 mL was collected from the tube and the peak fractions were confirmed by the real-time quantitative PCR as described above. The fractions containing vector particles were collected and applied to three rounds of dialysis against PBS using a Slide-A-Lyzer G2 dialysis cassette (MWCO 10,000; Thermo Fisher Scientific). The dialyzed solution was finally concentrated by centrifugation through a Vivaspin Turbo 4 membrane filter (MWCO 10,000; Sartorius) at 3,500 × *g* for 30-45 min at 4°C.

Viral genome titer was determined by quantitative PCR using a TaqMan system (Thermo Fisher Scientific). PCR amplification was performed using a StepOne real-time PCR system (Applied Biosystems) at 95°C, one cycle of 3 minutes, at 95°C 15 seconds, and 40 cycles at 60°C for 30 seconds for duplicate samples. Standard curves were prepared based on serial dilutions of the viral DNA control template from 4.0 × 10^4^ to 4.0 × 10^7^ genome copies/ml.

### Viral vector injection

*Drd2*-Cre rats were anesthetized with isoflurane (4% induction and 1.5% maintenance) and secured in a stereotaxic frame (SR-6R-HT, Narishige). Lenti-FLEX-EGFP/IR84a-2A-IR8a vector (4.36 × 10^12^ genome copies/ml for histological, behavioral, and biochemical experiments; 1.68 × 10^12^ genome copies/ml for *ex vivo* electrophysiological experiment), Lenti-FLEX-EGFP vector (4.97 × 10^13^ genome copies/ml for *ex vivo* electrophysiological experiment), or a 1:1 mixture of AAV2-EF1α-FLEX-EGFP/IR84a and AAV2-EF1α-FLEX-HA/IR8a (2.16 × 10^12^ genome copies/ml and 3.31 × 10^12^ genome copies/ml, respectively, for *in vivo* PET and *ex vivo* ARG experiments) were injected into the dorsal striatum through a glass microinjection capillary connected to a 10-μl gas-tight syringe (1801N, Hamilton) set in a microinfusion pump (ESP-32; Eicom). Injection was performed at a constant rate of 0.1 μl/min for 5 min according to the following anteroposterior (AP), mediolateral (ML), and dorsoventral (DV) coordinates from the bregma and dura (mm) of the rat brain (Paxinos and Watson, 2006): +0.15/±3.8/−4.4 (Site 1), +0.15/±3.8/−3.4 (Site 2), +0.15/±4.0/−4.4 (Site 3), +0.15/±4.0/−3.4 (Site 4) (bilateral) for histological experiments (Figure 3A); +0.15/±3.8/−4.4 (Site 1), +0.15/±3.8/−3.4 (Site 2), +0.15/±4.0/−4.4 (Site 3), +0.15/±4.0/−3.4 (Site 4), +0.15/±4.2/−4.4 (Site 5), +0.15/±4.2/−3.4 (Site 6), −0.35/±3.7/−4.7 (Site 7), −0.35/±3.7/−3.7 (Site 8), –0.35/±3.9−4.7 (Site 9), −0.35/±3.9/−3.7 (Site 10), −0.35/±4.1/−4.7 (Site 11), −0.35/±4.1/−3.7 (Site 12) for *ex vivo* electrophysiological experiment (bilateral, Figure 3B-D) and biochemical experiments (unilateral, random assignment of the viral vector injection to either the left or right hemisphere, Figure 4B-D); +1.44/2.0/−6.0 (Site 1), +1.44/2.0/−5.0 (Site 2), +1.44/3.0/−6.0 (Site 3), +1.44/3.0/−5.0 (Site 4), +0.6/2.5/−6.0 (Site 5), +0.6/2.5/−5.0 (Site 6), +0.6/3.5/−6.0 (Site 7), +0.6/3.5/−5.0 (Site 8) (unilateral) for *in vivo* PET and *ex vivo* ARG imaging (Figure 5B and C).

### Histology

Rats were anesthetized with sodium pentobarbital (100 mg/kg, ip.) and perfused transcardially with PBS, then fixed with 4% paraformaldehyde in 0.1 M phosphate buffer (pH 7.4). Fixed brain were cut with a vibratome (VT1000S, Leica), and then sections (50-μm thick) were incubated with 10% normal donkey serum, and then mixtures of a primary antibodies for Cre (mouse, 1:200; MAB3120, Sigma-Aldrich), A_2A_R (goat, 1:200; A2A-Go-Af700, Frontier Institute), and D1R (guinea pig, 1:200; D1R-GP-Af500, Frontier Institute) (for Figure 3—figure supplement 1), or GFP (chicken, 1:1,000; ab13970, Abcam), IR8a (guinea pig, 1:1,000; Abuin et al., 2011), and A_2A_R, or GFP, IR8a, and D1R (goat, 1:200, D1R-Go-Af1000, Frontier Institute) (for Figure 3A), or GFP and IR8a (for Figure 4—figure supplement 1A). Alexa Fluor 647-conjugated donkey anti-mouse antibody (A-31571, Thermo Fisher Scientific), Alexa Fluor 488-conjugated donkey anti-goat antibody (A-11055, Thermo Fisher Scientific), Cy3-conjugated donkey anti-guinea pig antibody (706-165-148, Jackson ImmunoResearch Laboratories), Alexa Fluor 488-conjugated donkey anti-chicken antibody (703-545-155, Jackson ImmunoResearch Laboratories), Alexa Fluor 647-conjugated donkey anti-goat antibody (1:200; A-21447, Thermo Fisher Scientific) were used as species-specific secondary antibodies for detecting Cre, A_2A_R, D1R, GFP, and A_2A_R, respectively (all dilutions were 1:200). Fluorescent images were obtained with a confocal laser-scanning microscope (A1, Nikon) or all-in-one fluorescence microscope (BZ-X810, Keyence) equipped with proper filter cube specifications.

For cell counts of triple-fluorescence histochemistries for GFP (IR84a), IR8a, and A_2A_R, and GFP, IR8a, and D1R in rats (Figure 3A), 4-5 sections through the striatum were prepared from each hemisphere and used for immunostaining. The number of immuno-positive cells in the region of interest was counted by using a computer-assisted imaging program (Photoshop 2021; Adobe). Number of cells of each type from 5-7 visual fields of the same hemisphere (each was 200 × 200 μm, a total of 50-80 cells were identified) was counted, and the means of 4 hemispheres were shown in Supplementary Table 1. The IR84a/IR8a double-expression efficiency in the A_2A_R– and D1R-positive neurons was calculated by dividing the number of IR84a/IR8a double-positive neurons by the number of triple-positive neurons. The IR84a/IR8a double-expression specificity was calculated by dividing the number of triple-positive neurons by the number of IR84a/IR8a double-positive.

### Electrophysiology

*Ex vivo* electrophysiology with brain slices was performed as described in Fukabori et al. (2020). Briefly, animals were anesthetized with 1.5% isoflurane. Coronal brain slices containing the LC (in mice, for Figure 2—figure supplement 1A-F) or striatum (in rats, for Figure 3B-D) were cut (300 μm thick) using a microslicer (PRO7, Dosaka) in ice-cold oxygenated cutting Krebs solution of the following composition (mM): choline chloride, 120, KCl, 2.5, NaHCO_3_, 26, NaH_2_PO_4_, 1.25, D-glucose, 15, ascorbic acid, 1.3, CaCl_2_, 0.5, and MgCl_2_, 7. The slices were then transferred to a holding chamber containing a standard Krebs solution of the following composition (mM): NaCl, 124, KCl, 3, NaHCO_3_, 26, NaH_2_PO_4_, 1, CaCl_2_, 2.4, MgCl_2_, 1.2, and D-glucose, 10 (pH 7.4) when bubbled with 95% O_2_-5% CO_2_. Slices were incubated in the holding chamber at room temperature (21°-26°C) for at least 1 hour before recording. Neurons were visualized with a 60× water immersion objective attached to an upright microscope (BX50WI, Olympus Optics), and fluorescence was visualized using the appropriate fluorescence filter. Patch pipettes were made from standard-walled borosilicate glass capillaries (Harvard Apparatus). For the recording of membrane potentials, a K-gluconate-based internal solution of the following composition (mM) was used: K-gluconate, 120, NaCl, 6, CaCl_2_, 5, MgCl_2_, 2, K-EGTA, 0.2, K-HEPES, 10, Mg-ATP, 2, and Na-GTP, 0.3 (pH adjusted to 7.4 with 1 M KOH). Whole-cell recordings were made from LC neurons with fluorescence using a patch-clamp amplifier (Axopatch 200B, Molecular Devices). The effects of drugs (*S*)-PhPrM and (*R*)-PhPrM for mice LC neurons (Figure 2—figure supplement 1A-F), PhAcM for rats striatal neurons (Figure 3B-D) on the membrane potential were assessed after they had reached a steady state (starting point), and the mean firing frequency and membrane potential were calculated during a 30-s test period before the drug application (pre) and after the starting point (post).

*In vivo* extracellular single-unit recording with the Tg mice (for Figure 1B, Figure 2B and C) was carried out as previously described (Fukabori et al., 2020). Briefly, mice were anesthetized with 1.5% isoflurane and placed in the stereotaxic frame (SR-5M, Narishige) with ear bars and a mouth-and-nose clamp. Anesthesia was maintained with 0.5-1.0% isoflurane based on the electroencephalogram monitoring with body temperature at 37-38°C using a heating pad. The scalp was opened, and a hole was drilled in the skull above the LC with the coordinates (in mm) AP −1.2 and ML ±0.9 from lambda according to the mouse stereotaxic atlas (Paxinos and Franklin, 2008). Two skull screws were placed over the occipital bone, and another skull screw was placed on the frontal bone. One of the two screws over the occipital bone was used as a reference for the recording. The firing activity of LC neurons was recorded using a recording electrode (tip diameter: 2-3 μm, impedance: 15-20 MΩ, Harvard Apparatus), which was lowered into the LC, dorsoventral (DV) −2.2 to −3.5 mm from the dura, and recording was performed using Spike2 (Cambridge Electronic Design) at a sampling rate of 20 kHz. Solution of PhAcM (20 mg/kg), (*S*)-PhPrM (20 mg/kg) or vehicle was administered in the lateral tail vein at 5 ml/kg. The baseline firing rate (pre) was recorded during a 180-s period immediately before drug administration. Ten mins after the drug administration, the effects were assessed for 10 min (post for PhAcM, Figure 1B) or 90 min (post for (*S*)-PhPrM, Figure 2B-C).

After the electrophysiological experiments, the recording sites were marked by iontophoretic injection of 2% pontamine sky blue. The mice were deeply anesthetized with sodium pentobarbital and perfused transcardially with PBS, followed by 10% formalin. Brain sections were stained with neutral red for the verification of recording sites. The post-mortem histological analysis verified deposits of recording sites in the LC (Figure 1—figure supplement 2A, Figure 2—figure supplement 3A).

### Microdialysis

To monitor the change in the NE_[ext]_ in the ACC, mice were anesthetized with 1.5% isoflurane and underwent stereotaxic surgery for a dialysis probe aimed at unilateral ACC with 25-gauge guide cannula (AG-X series, Eicom). A group of mice (for Figure 2—figure supplement 1G and H) also underwent stereotaxic surgery for drug microinjection into the ipsilateral LC with a 30-gauge guide cannula (AG-X(T) series, Eicom). The coordinates (mm) from bregma or dura were AP +0.7, ML ±0.3, and DV −0.4 for the ACC dialysis probe and AP −0.75, ML ±0.65, and DV −2.5 for the LC microinjection, according to the mouse stereotaxic atlas (Paxinos and Franklin, 2008). Two or three days later, the stylet in the cannula was replaced with an active membrane dialysis probe (1.0 mm in length, 0.22 mm in outer diameter, FX-I series, Eicom) that was connected to a 2,500 μl syringe filled with artificial cerebrospinal fluid (aCSF) of the following composition (mM): NaCl, 148, KCl, 4.0, MgCl_2_, 0.85, and CaCl_2,_ 1.2. For the intra-LC injection, the stylet in the LC cannula was replaced with a 35-gauge internal cannula (1 mm beyond the tip of the implanted guide cannula, AMI-X(T) series, Eicom) connected to a 10 μl Hamilton syringe. aCSF was pumped through the probe at a rate of 1.0 μl/min for 2 h, and then dialysate samples were collected in sampling vials every 30 min using a refrigerated fraction collector (EFC-82, Eicom). Each sampling vial was pre-loaded with 10 μl of 20 mM phosphate buffer, including 25 mM EDTA-2Na and 0.5 mM of ascorbic acid (pH 3.5) as antioxidants. Three baseline samples were collected to measure a tonic level of NE. This was followed by tail vein injection of PhAc (Figure 1—figure supplement 1A), PhAcM (Figure 1C), or (*S*)-PhPrM (Figure 2D), or intracranial LC microinjection (at a flow rate of 0.05 μl/min for 5 min) of 0.6% (*S*)-PhPr or 0.6% (*R*)-PhPr (Figure 2—figure supplement 1G). Four samples were collected thereafter to assess the time course for change in NE level after these interventions. The amount of NE in each fraction was determined by a high-performance liquid chromatography (HPLC) system (CA-50DS, 2.1 mm × 150 mm, Eicom, with the mobile phase containing 5% methanol in 100 mM sodium phosphate buffer, pH 6.0) equipped with an electrochemical detector (ECD-300, Eicom). Results are expressed as a percentage of baseline concentration (analyte concentration × 100/mean of the three baseline samples).

To monitor the change in the GABA_[ext]_ in the GPe, the viral vector-treated Drd2-Cre rats that had been subjected to behavioral experiment were anesthetized with 1.5% isoflurane and underwent stereotaxic surgery for a dialysis probe aimed at bilateral GPe with 25-gauge guide cannula. According to the rat stereotaxic atlas, the coordinates (mm) from bregma or dura were AP −0.9, ML ±3.0, and DV −5.1 (Paxinos and Watson, 2006). Seven to 10 days later, the stylet in the cannula was replaced with an active membrane dialysis probe, and aCSF was perfused through the probe at a rate of 1.0 μl/min for 2 h, and then dialysate samples were collected sampling vials every 30 min using a refrigerated fraction collector. Each dialysate sample was reacted with 10 μl of 20 mM o-phthalaldehyde, including 0.2% 2-mercaptoethanol, for 2.5 min and then injected into HPLC at a volume of 30 μl (SC-50DS, 2.1 mm × 150 mm, Eicom, with the mobile phase containing 50% methanol in 50 mM sodium phosphate buffer, pH 6.0) equipped with an electrochemical detector. Three baseline samples were collected to measure GABA tone. This was followed by a tail vein injection of 10 mg/kg PhAcM (Figure 4C) or 20 mg/kg (*S*)-PhPrM (Figure 4D). Four samples were collected thereafter to assess the time course for change in GABA levels after these interventions. Results are expressed as a percentage of baseline concentration (analyte concentration × 100/mean of the three baseline samples).

After the microdialysis experiments, animals were deeply anesthetized with sodium pentobarbital and perfused transcardially with PBS, followed by 10% formalin. Brain sections were stained with cresyl violet for the verification of placement sites. The post-mortem histological analysis verified deposits of dialysis probe sites in the ACC of mice (Figure 1—figure supplement 1C, Figure 1—figure supplement 2B, Figure 2—figure supplement 1H, Figure 2—figure supplement 3B) and the GPe of rats (Figure 4—figure supplement 1B) as well as microinjection sites in the LC of mice (Figure 2—figure supplement 1H).

### Behavioral analysis

The rotational behavior of rats was examined in a transparent, acrylic plastic cylinder (30 cm in diameter, 55 cm in height) with a hemispheric bottom made of as the apparatus. A two-day test was conducted after 14 days of recovery from the stereotaxic surgery for viral vector administration, during which animals were handled and acclimated to the apparatus. On day 1 of the two-day test, rats were given 10 mg/kg PhAcM and 15 mg/kg cocaine i.p. consecutively at 10-min interval and then collared and placed in the apparatus for 60 min to monitor their rotations. The collar was connected, via a stainless wire, to a rotary encoder (E6A2-CWZ3C, Omron), which determines clockwise and anti-clockwise rotations of animals. The rotation information was transferred to a personal computer via an interface (USB-6501, National Instruments) and recorded using a Matlab simulink-based application (Matlab R2019b, Mathworks). The procedure for day 2 of the test was performed 48 hr after day 1 similarly, except that a vehicle was administered instead of PhAcM. Rotational behavior recorded during the 60-min test sessions was analyzed for ipsiversive and contraversive ones of the viral vector-treated hemisphere every 5-min bins. The range of expression of IR84a (GFP)/IR8a double-positive cells in the rat striatum was visualized by the post-mortem histological analysis (Figure 4—figure supplement 1A).

### PET imaging and *ex vivo* ARG

Rats were placed under isoflurane anesthesia with a plastic catheter (26G) in their tail vein for PET tracer administration. The rats were then set in an animal PET scanner (microPET Focus-220, Siemens Medical Solutions, Knoxville, TN, USA) for a head scan. A bolus injection of (*S*)-[^11^C]PhPrM was performed at 68.7-75.2 MBq/0.3 ml in physiological saline over 10 s. After dosing, saline was flushed with 0.5 ml of saline. The emission scan was done at the same time as the dose for 60 min. During the PET scanning, the animal was kept under isoflurane anesthesia (2.2-3.0%). The acquired emission data reconstructed the PET images using a filtered back-projection algorithm with no scattered and attenuation corrections.

For *ex vivo* ARG, the brain tissue of the rats used for PET imaging was removed immediately after euthanasia under deep anesthesia. Brains were sectioned coronally 1 mm thick using a brain slicer under ice-cold conditions, and brain slices were placed in a chamber with plastic film attached and exposed to an imaging plate (IP, MS-2040, Fuji Film, Japan) for 40 min. The exposed IP was taken with an image analyzer (FLA-7000IR, Fuji Film). The photographs of brain slices were also acquired with a digital scanner as an anatomical reference for brain regions. To quantify ^11^C radioactivity, Multi Gage software (Fuji Film), the regions of interests in the vector-injected and opposite control sides were drawn on ARG images.

### Quantification and statistical analyses

Although no statistical methods were used to predetermine the sample size for each measure, we employed similar sample sizes to those reported in previous publications from our labs, which have been generally accepted in the field. In each test, we conducted an experiment in two to five cohorts, each designed to include all experimental groups. All statistical analyses were two-tailed and conducted using SPSS ver. 25 (IBM). The reliability of the results was assessed against a type I error (α) of 0.05. For significant main effects identified in one– and two-way analyses of variance (ANOVAs). For significant interactions revealed in two-way ANOVAs, t-tests with Bonferroni correction were used post hoc. Besides, all effect sizes were reported as partial η^2^ for main effects and interaction in ANOVAs and as r for pot hoc paired comparisons. In principle, the main text describes up to the main effects and interaction of ANOVAs and subsequent tests are reported in the corresponding figure legends. One star and daggers (*, †) represent a p-value < 0.05 in the figures.

## Data availability

The datasets generated and analyzed in this study are deposited on Mendeley data. DOI: 10.17632/b47j739jb7.1

## Contact for resources and reagents sharing

Further information and requests for resources and reagents should be directed to and will be fulfilled by the Lead Contact, Kazuto Kobayashi.

## Article and author information

### Author details

**Yoshio Iguchi**

Department of Molecular Genetics, Institute of Biomedical Sciences, Fukushima Medical University School of Medicine, Fukushima, Japan

**Contribution:** Conceptualization, Data curation, Software, Formal analysis, Validation, Investigation, Visualization, Methodology, Writing – original draft, Writing – review and editing, Funding acquisition

**Competing interests:** No competing interests declared

**ORCID:** https://orcid.org/0000-0001-8240-3345

**Ryoji Fukabori**

Department of Molecular Genetics, Institute of Biomedical Sciences, Fukushima Medical University School of Medicine, Fukushima, Japan

**Contribution:** Conceptualization, Resources, Validation, Investigation, Methodology, Writing – review and editing

**Competing interests:** No competing interests declared

**ORCID:** https://orcid.org/0000-0001-7534-8237

**Shigeki Kato**

Department of Molecular Genetics, Institute of Biomedical Sciences, Fukushima Medical University School of Medicine, Fukushima, Japan

**Contribution:** Resources, Validation, Investigation, Methodology, Writing – original draft, Writing – review and editing

**Competing interests:** No competing interests declared

**ORCID:** https://orcid.org/0000-0002-8792-7591

**Kazumi Takahashi**

Department of Systems Neuroscience, Fukushima Medical University School of Medicine, Fukushima Japan

**Contribution:** Data curation, Formal analysis, Validation, Investigation, Visualization, Methodology, Writing – original draft, Writing – review and editing

**Competing interests:** No competing interests declared

**ORCID:** https://orcid.org/0000-0003-4015-8016

**Satoshi Eifuku**

Department of Systems Neuroscience, Fukushima Medical University School of Medicine, Fukushima Japan

**Contribution:** Validation, Writing – review and editing, Supervision

**Competing interests:** No competing interests declared

**ORCID:** https://orcid.org/0000-0003-0032-1720

**Yuko Maejima**

Department of Bioregulation and Pharmacological Medicine, Fukushima Medical University School of Medicine, Fukushima Japan

**Contribution:** Validation, Investigation, Visualization, Methodology, Writing – original draft, Writing – review and editing

**Competing interests:** No competing interests declared

**ORCID**:

**Kenju Shimomura**

Department of Bioregulation and Pharmacological Medicine, Fukushima Medical University School of Medicine, Fukushima Japan

**Contribution:** Data curation, Formal analysis, Validation, Investigation, Visualization, Methodology, Writing – original draft, Writing – review and editing, Supervision

**Competing interests:** No competing interests declared

**ORCID**:

**Hiroshi Mizuma**

Laboratory for Pathophysiological and Health Science, RIKEN Center for Biosystems Dynamics Research, Kobe, Japan

Department of Functional Brain Imaging, Institute for Quantum Medical Science, National Institutes for Quantum Science and Technology, Chiba, Japan

**Contribution:** Resources, Data curation, Formal analysis, Validation, Investigation, Visualization, Methodology, Writing – original draft, Writing – review and editing

**Competing interests:** No competing interests declared

**ORCID:** https://orcid.org/0000-0001-8970-9486

**Aya Mawatari**

Laboratory for Labeling Chemistry, RIKEN Center for Biosystems Dynamics Research, Kobe, Japan

**Contribution:** Resources, Validation, Investigation, Methodology, Writing – original draft, Writing – review and editing

**Competing interests:** No competing interests declared

**ORCID**:

**Hisashi Doi**

Laboratory for Labeling Chemistry, RIKEN Center for Biosystems Dynamics Research, Kobe, Japan Research Institute for Drug Discovery Science, Collaborative Creation Research Center, Organization for Research Promotion, Osaka Metropolitan University, Osaka, Japan

**Contribution:** Resources, Validation, Investigation, Methodology, Writing – original draft, Writing – review and editing

**Competing interests:** No competing interests declared

**ORCID:** https://orcid.org/0000-0001-9778-5841

**Yilong Cui**

Laboratory for Biofunction Dynamics Imaging, RIKEN Center for Biosystems Dynamics Research, Kobe, Japan

**Contribution:** Conceptualization, Data curation, Formal analysis, Validation, Investigation, Visualization, Methodology, Writing – original draft, Writing – review and editing, Supervision

**Competing interests:** No competing interests declared

**ORCID:** https://orcid.org/0000-0002-8302-1899

**Hirotaka Onoe**

Human Brain Research Center, Kyoto University Graduate School of Medicine, Kyoto, Japan

**Contribution:** Conceptualization, Validation, Methodology, Writing – review and editing, Supervision

**Competing interests:** No competing interests declared

**ORCID**:

**Keigo Hikishima**

Medical Devices Research Group, Health and Medical Research Institute, National Institute of Advanced Industrial Science and Technology (AIST), Tsukuba, Japan

**Contribution:** Validation, Methodology, Writing – review and editing

**Competing interests:** No competing interests declared

**ORCID:** https://orcid.org/0000-0003-1133-1297

**Makoto Osanai**

Department of Medical Physics and Engineering, Division of Health Sciences, Osaka University Graduate School of Medicine, Suita, Japan

**Contribution:** Validation, Methodology, Writing – review and editing, Supervision

**Competing interests:** No competing interests declared

**ORCID:** https://orcid.org/0000-0002-3572-8195

**Takuma Nishijo**

Department of Pharmacology, Jikei University School of Medicine, Tokyo, Japan Department of Molecular Neurobiology, Institute for Developmental Research, Aichi Developmental Disability Center, Kasugai, Japan

**Contribution:** Validation, Investigation, Visualization, Methodology, Writing – original draft, Writing – review and editing

**Competing interests:** No competing interests declared

**ORCID:** https://orcid.org/0000-0003-0533-0132

**Toshihiko Momiyama**

Department of Pharmacology, Jikei University School of Medicine, Tokyo, Japan **Contribution:** Data curation, Formal analysis, Validation, Investigation, Visualization, Methodology, Writing – original draft, Writing – review and editing, Supervision **Competing interests:** No competing interests declared

**ORCID:**

**Richard Benton**

Center for Integrative Genomics, Faculty of Biology and Medicine, University of Lausanne, Lausanne, Switzerland

**Contribution:** Resources, Validation, Methodology, Writing – original draft, Writing – review and editing, Supervision

**Competing interests:** No competing interests declared

**ORCID:** https://orcid.org/0000-0003-4305-8301

**Kazuto Kobayashi**

Department of Molecular Genetics, Institute of Biomedical Sciences, Fukushima Medical University School of Medicine, Fukushima, Japan

**Contribution:** Conceptualization, Resources, Data curation, Validation, Visualization, Methodology, Writing – original draft, Writing – review and editing, Funding acquisition, Supervision

**Competing interests:** No competing interests declared

**ORCID:** https://orcid.org/0000-0002-7617-2939

## Funding

**Ministry of Education, Science, Sports (MEXT), Japan, Scientific Research on Innovative Areas “Adaptive Circuit Shift” (26112002)**

Kazuto Kobayashi

**Ministry of Education, Science, Sports (MEXT), Japan, Transformative Research Areas “Adaptive Circuit Census” (21H05244)**

Kazuto Kobayashi

**Japan Society for the Promotion of Science (JSPS), Challenging Research (Exploratory) (22K19363)**

Kazuto Kobayashi

**Takeda Science Foundation (2021), Japan**

Kazuto Kobayashi

**European Research Council Consolidator Grant (Grant 615094)**

Richard Benton

**Human Frontier Science Program Young Investigator Award (Grant RGY0073/2011)**

Richard Benton

**Swiss National Science Foundation Nano-Tera Envirobot Project (20NA21_143082)**

Richard Benton

**Japan Society for the Promotion of Science (JSPS), Scientific Research (C) (19K03381)**

Yoshio Iguchi

**Takeda Science Foundation (2020), Japan**

Yoshio Iguchi

The funders had no role in study design, data collection, interpretation, or the decision to submit the work for publication.

## Acknowledgments

We are grateful to M. Kikuchi and N. Sato for their maintenance and management of laboratory animals, Y. Nakazato for her viral vector preparation, and H. Hashimoto and S. Chitose for their technical support.

## Supplementary figure legends

**Figure 1—figure supplement 1. Peripheral administration of PhAc does not stimulate the IR84a/IR8a-expressing neurons in the brain.** (**A**) Plot of changes in the NE_[ext]_ in the ACC of the TH-IR84a/IR8a (Tg) mice before and after tail vein injection with vehicle or PhAc (10 or 30 mg/kg). NE_[ext]_ is expressed as a percentage of the average baseline levels of each mouse. Arrow indicates the timing of drug injection. NE_[ext]_ did not show any significant changes among drug administrations (ns = 4 for vehicle and PhAc 10 mg/kg, n = 6 for PhAc 30 mg/kg); two-way ANOVA, dose effect: F_(2, 11)_ = 0.28, p = 0.762, partial η^2^ = 0.05, fraction effect: F_(6, 66)_ = 0.49, p = 0.813, partial η^2^ = 0.04, interaction: F_(12, 66)_ = 0.78, p = 0.671, partial η^2^ = 0.12. (**B**) Plot of changes in NE_[ext]_ in the ACC of the Tg mice before and after reverse dialysis with PhAc (3 – 3,000 μM). Horizontal bar indicates the period of PhAc perfusion. A perfusion of different concentrations of PhAc into the dialysis probe elevated NE_[ext]_ in a dose-dependent manner (n = 3 for each condition); two-way ANOVA, concentration effect: F_(3, 8)_ = 27.14, p < 0.001, partial η^2^ = 0.91, fraction effect: F_(3, 24)_ = 7.38, p = 0.001, partial η^2^ = 0.48, interaction: F_(9, 24)_ = 2.38, p = 0.044, partial η^2^ = 0.47; NE_[ext]_ following the 3,000-μM PhAc perfusion was significantly higher than those following the *3– and ^†^30-μM PhAc at the 30-min fraction, ts_(32)_ = 4.41, 3.18, p < 0.001, p = 0.020, rs = 0.62, 0.49, respectively, and that following the *3-μM PhAc at the 60-min fraction, t_(32)_ = 3.24, p = 0.017, r = 0.50, whereas NE_[ext]_ following the 300-μM PhAc perfusion was significantly higher than that following the *3 μM PhAc at the 30-min fraction, t_(32)_ = 3.59, p = 0.007, r = 0.54, and those following the *3– and ^†^30-μM PhAc at the 60-min fraction, ts_(32)_ = 4.33, 3.35, p < 0.001, p = 0.012, rs = 0.61, 0.51, respectively. Data are presented as mean ± SEM. (**C**) Site mapping of the dialysis probes in the ACC of the Tg mice. The anteroposterior coordinates (mm) are shown. Scale bar: 1 mm.

**Figure 1—figure supplement 2. Enhancement of the BBB permeability of PhAc by methyl esterification.** (**A)** Site mapping of the tips of glass electrode in the LC of the Tg mice (for Figure 1B). (**B**) Site mapping of the dialysis probes in the ACC of the Tg and non-Tg mice (for Figure 1C). The anteroposterior coordinates (mm) are shown. Scale bar: 1 mm.

**Figure 2—figure supplement 1. The *S*-isomer of PhPr stimulates IR84a/IR8a-expressing neurons.** (**A**) Representative whole-cell recordings of the LC neurons of the control (TH-GFP) and Tg (TH-IR84a/IR8a) mice in response to (*S*)-PhPr. (**B-C**) Plot of changes in the membrane potential (**B**) and firing rate (**C**) with addition of (*S*)-PhPr (0.1% w/v). Each white dot in the boxplot indicate mean for the condition. The amount of difference in the membrane potential between pre– and post-(*S*)-PhPr (0.1%) bath application (Δ mV) of the LC-NE neurons in the Tg mice was significantly greater than that of the LC-NE neurons in the control mice (n = 4 for each group); one-way ANOVA, genotype effect: F(1, 6) = 13.65, p = 0.010, partial η^2^ = 0.69. The normalized firing rate was also significantly elevated by the (*S*)-PhPr application in the Tg mice but not in the control mice; one-way ANOVA for the Tg mice, genotype effect: F(1, 3) = 109.13, p = 0.002, partial η^2^ = 0.97; for the control mice, F(1, 3) = 0.05, p = 0.836, partial η^2^ = 0.02. **(D**) Representative whole-cell recordings of the LC neurons of the control and Tg mice in response to (*R*)-PhPr. (**E-F**) Plot of changes in the membrane potential (E) and firing rate (F) with addition of (*R*)-PhPr (0.1% w/v). Δ mV by (*R*)-PhPr (0.1%) was almost zero for both control and Tg mice and no significant difference was found between the two conditions (n = 4 for each group); one-way ANOVA, genotype effect: F(1, 6) = 1.64, p = 0.247, partial η^2^ = 0.21. (*R*)-PhPr had no significant impacts on the normalized firing rate of not only control but also Tg mice; one-way ANOVA for the control mice, genotype effect: F(1, 3) = 0.94, p = 0.403, partial η^2^ = 0.24; for the Tg mice, F(1, 3) = 2.64, p = 0.203, partial η^2^ = 0.47. (**G**) Plot of changes in the extracellular NE level (NE_[ext]_) in the ACC area of the Tg and non-Tg mice before and after microinjection of the (*S*)– and (*R*)-PhPr (0.6% w/v) into the LC. (*S*)-PhPr resulted in a slightly delayed but marked increase in NE_[ext]_ in the Tg mice (ns = 4 for the two (*S*)-PhPr groups, n = 3 for (*R*)-PhPr group); two-way ANOVA, group effect: F(2, 8) = 26.42, p < 0.001, partial η^2^ = 0.89, fraction effect: F(6, 48) = 1.21, p = 0.318, partial η^2^ = 0.13, interaction: F(12, 48) = 2.01, p = 0.044, partial η^2^ = 0.33; NE_[ext]_ of (*S*)-PhPr in the Tg mice was significantly higher than that of *(*R*)-PhPr in the Tg mice and ^†^(*S*)-PhPr in the non-Tg mice at the 60-min fractions, ts(56) = 4.82, 5.19, ps < 0.001, rs = 0.54, 0.57, respectively. Data are presented as mean ± SEM. (**H**) Site mapping of the dialysis probes in the ACC and the injection cannula tips near the LC of the Tg and non-Tg mice (for **G**). The anteroposterior coordinates (mm) are shown. Scale bar: 1 mm.

**Figure 2—figure supplement 2. TLC analysis of the tissue sample dissected from the LC area of the Tg mouse brain after (*S*)-[^11^C]PhPrM treatment**. The Tg mouse was intravenously injected with (*S*)-[^11^C]PhPrM (42 MBq/0.1 ml), and its brain was removed 30 min later to prepare tissue suspension of the LC region for the TLC analysis. The sample and (*S*)-[^11^C]PhPrM were examined using a radio-TLC scanner (left photo), while the standard compounds were visualized using standard methods (UV light, right photo). Retention factor (Rf) values for (*S*)-[^11^C]PhPrM and the LC tissue suspension were 0.36 and 0.60, respectively, and Rf values for (*S*)-PhPrM and (*S*)-PhPr were 0.38 and 0.64, respectively. The data indicate that the value obtained from the tissue suspension was similar to that from the standard of (*S*)-PhPr, suggesting the conversion of (*S*)-[^11^C]PhPrM in the Tg mouse brain.

**Figure 2—figure supplement 3. Placement sites of glass electrodes for in vivo electrophysiology and dialysis probes for microdialysis analysis.** (**A)** Site mapping of the tips of glass electrodes in the LC of the Tg mice (for Figures 2C and D). (**B**) Site mapping of the dialysis probes in the ACC of the Tg and non-Tg mice (for Figure 2E). The anteroposterior coordinates (mm) are shown. Scale bar: 1 mm.

**Figure 3—figure supplement 1. Specificity of the Cre recombinase expression in iSPNs of the striatum of the *Drd2*-Cre rats.** Triple immunohistochemistry of the section through the striatum of the *Drd2*-Cre rats for Cre, A_2A_R, and D1R established that the Cre transgene was highly and specifically expressed in iSPNs (A_2A_R-positive) but was absent in dSPNs (D1R-positive). + Cre-positive cell, – Cre-negative cell. Scale bar: 25 μm.

**Figure 4—figure supplement 1. Expression patterns of transgenes and placement sites dialysis probes for microdialysis analysis.** (**A)** Range of IR84a/IR8a expression in the dorsal striatum of the *Dr2d*-Cre rats (for Figure 4B). (**B**) Site mapping of the dialysis probes in the GPe of the *Dr2d*-Cre rats (for Figure 4C and D). The anteroposterior coordinates (mm) are shown. Scale bar: 1 mm.

## Supplementary Table

**Supplementary Table 1.** Count of various types of immuno-positive neurons in the viral vector-treated striatum of the *Drd2*-Cre rats

## Notes

https://doi.org/10.17632/b47j739jb7.1

### Competing Interest Statement

The authors have declared no competing interest.

