## Supplementary figures and images for "Chemogenetic activation of target neurons expressing insect Ionotropic Receptors in the mammalian central nervous system by systemic administration of ligand precursors"

### Supplemental Data 1

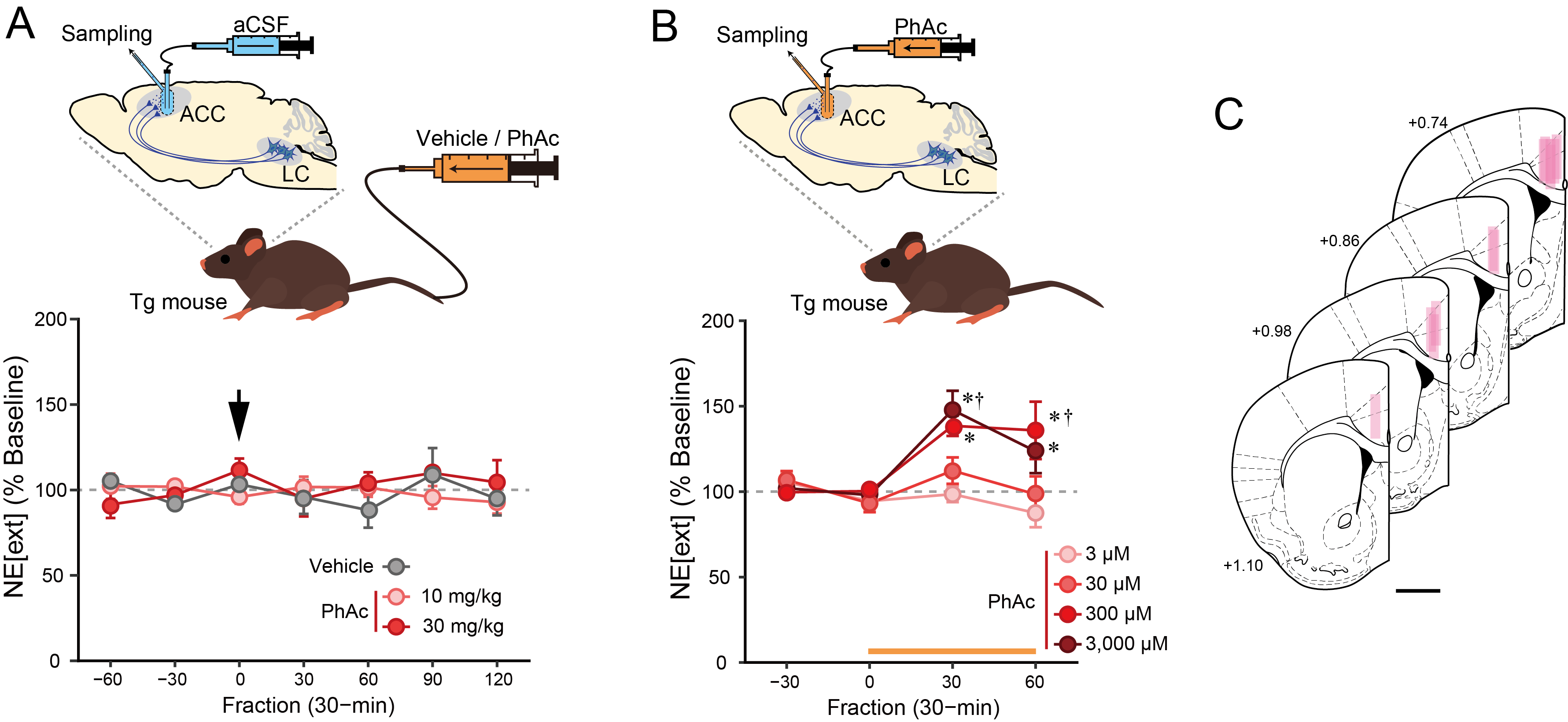

### Supplemental Data 2

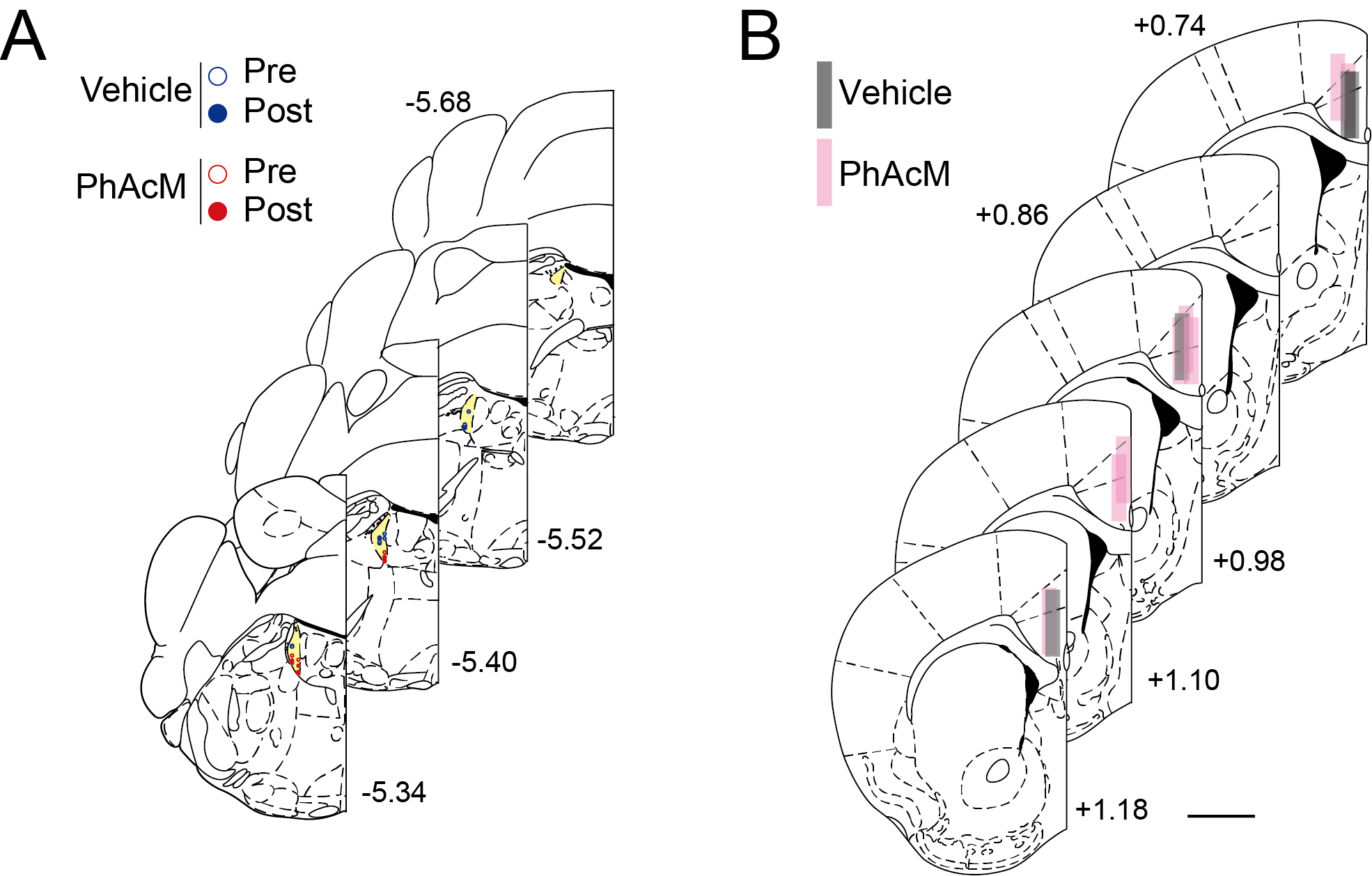

### Supplemental Data 3

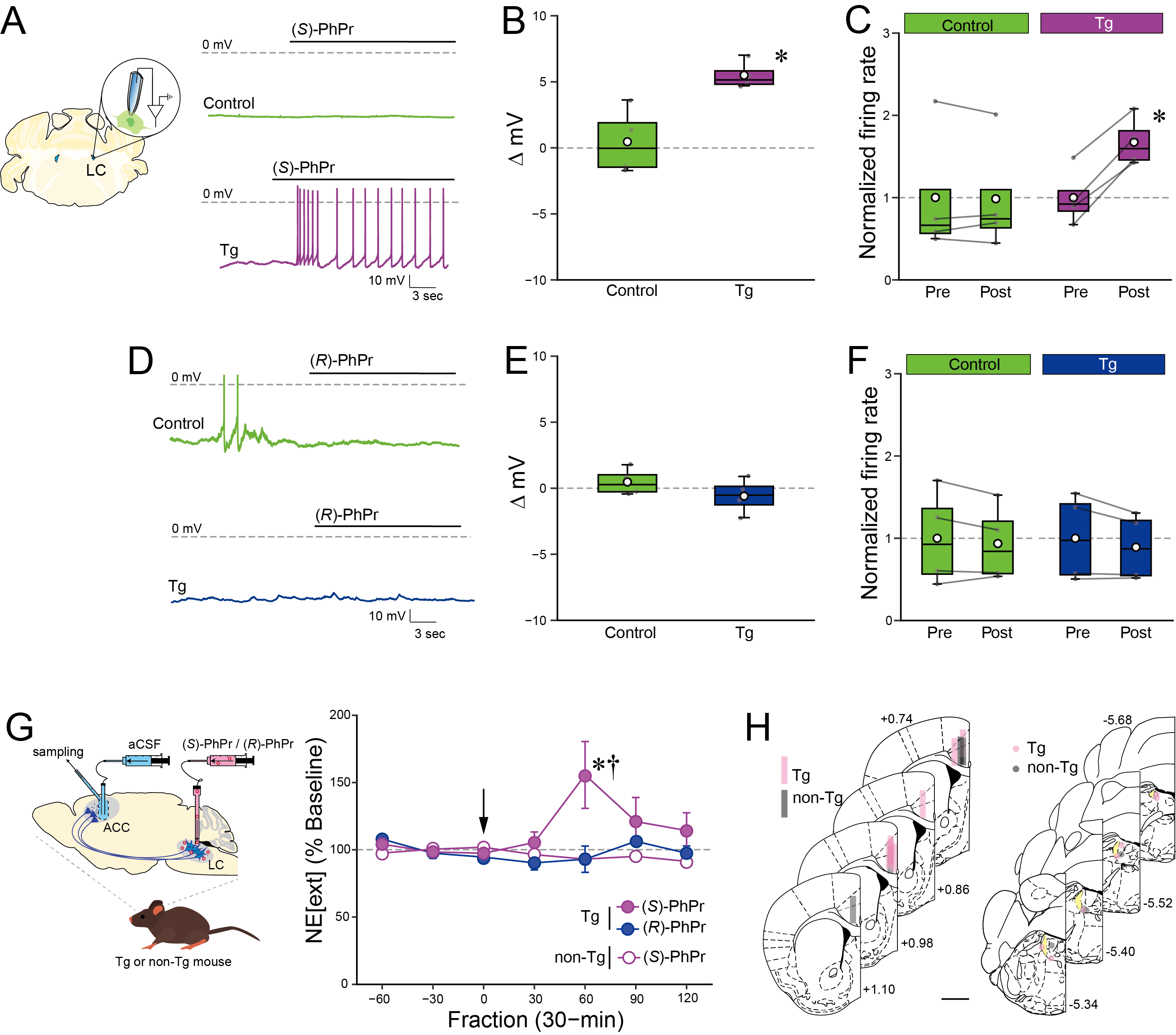

### Supplemental Data 4

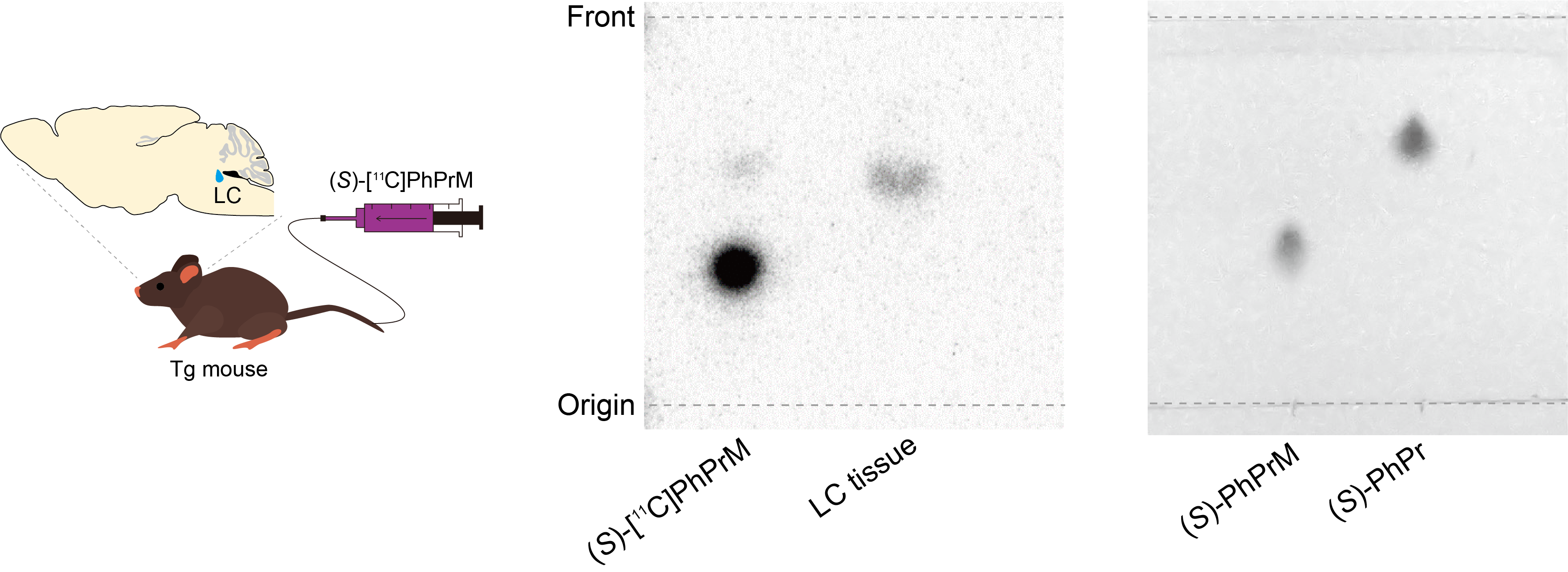

### Supplemental Data 5

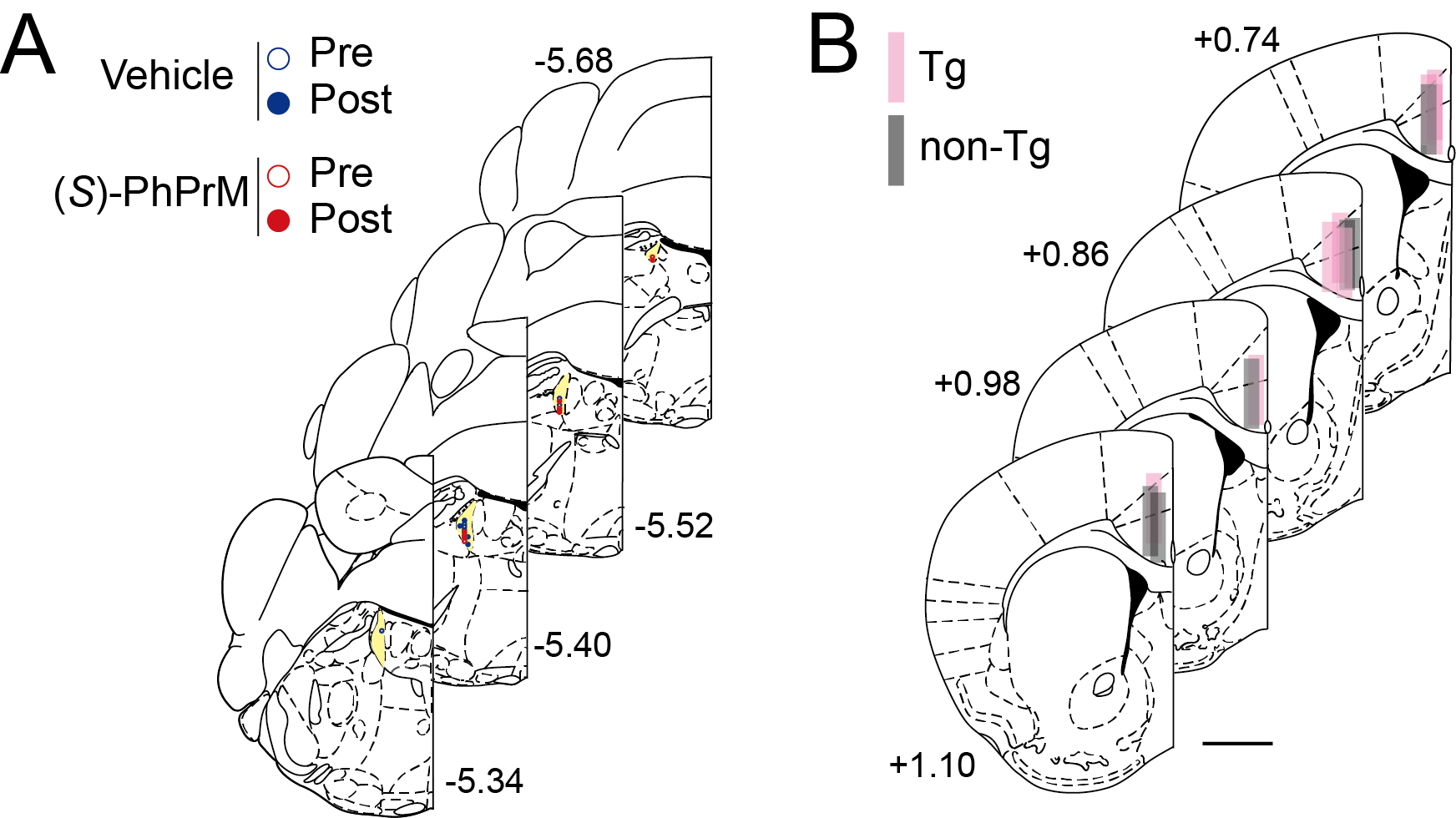

### Supplemental Data 6

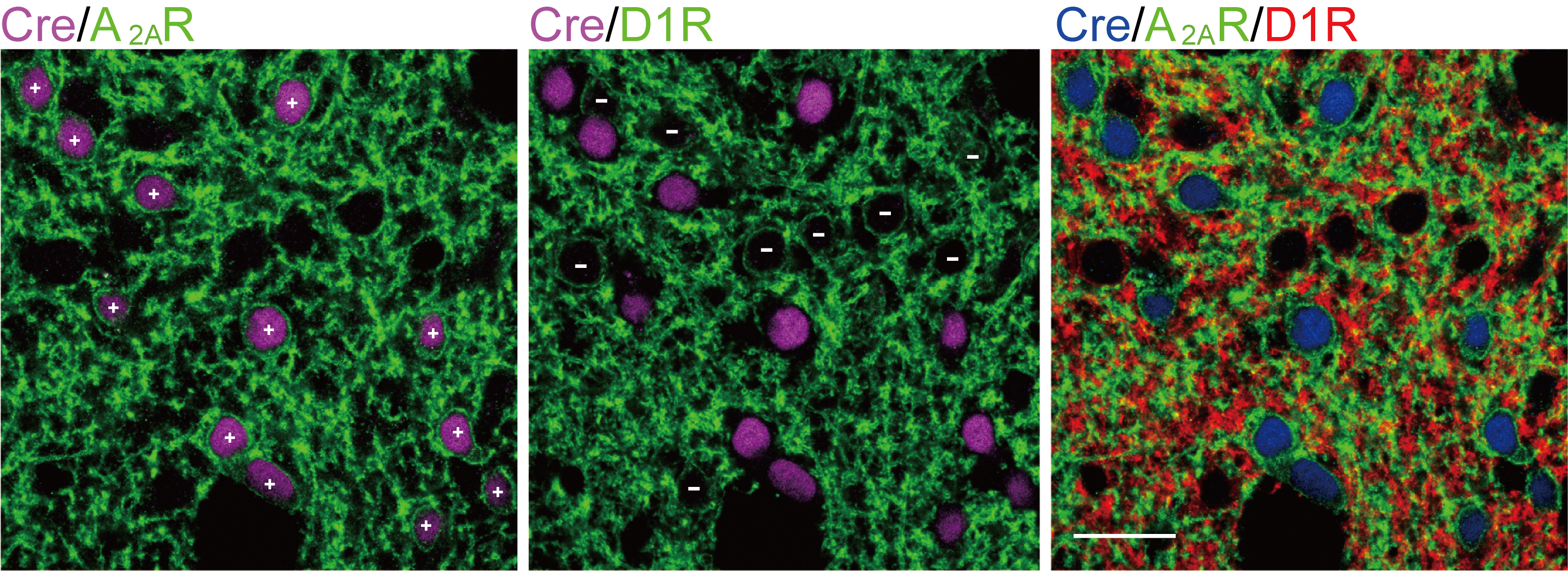

### Supplemental Data 7

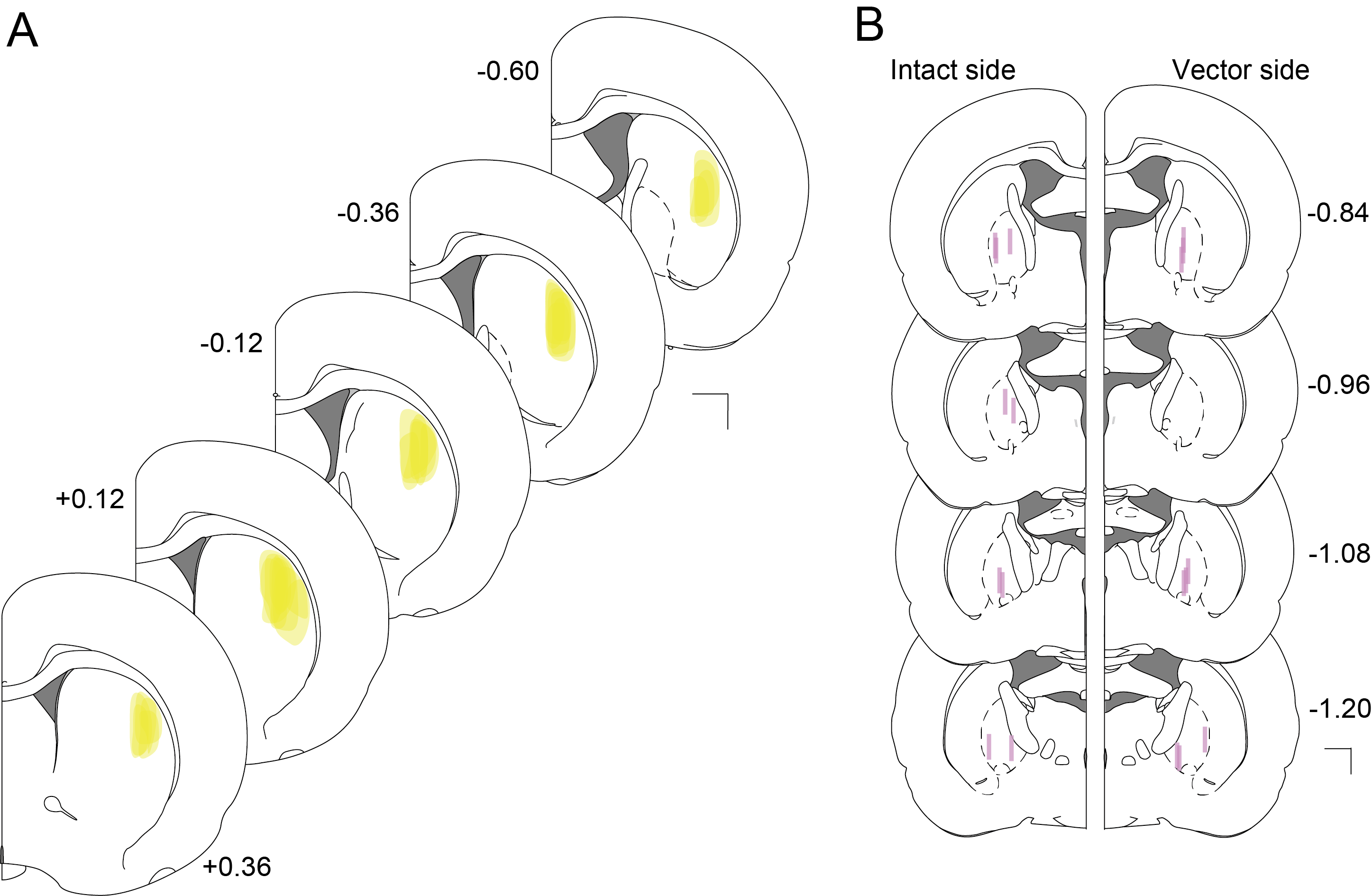
