## Supplementary Table 1 for "Chemogenetic activation of target neurons expressing insect Ionotropic Receptors in the mammalian central nervous system by systemic administration of ligand precursors"

**Supplementary Table 1. Count of various types of immuno-positive neurons in the viral vector-treated striatum of the *Drd2*-Cre rats**

|  | Section | |
| --- | --- | --- |
| Cell classification | Incubated w/ anti- A_2A_R-antibody | Incubated w/ anti-D1R-antibody |
| GFP (IR84a)^+^ | 65.50 ± 4.94 | 65.00 ± 5.80 |
| IR8a^+^ | 71.00 ± 4.34 | 68.50 ± 5.55 |
| GPF^+^ & IR8a^+^ | 63.00 ± 3.58 | 62.25 ± 6.12 |
| A_2A_R^+^ | 68.25 ± 5.22 | NA |
| D1R^+^ | NA | 52.50 ± 3.77 |
| GFP^+^ & A_2A_R^+^ | 51.50 ± 4.29 | NA |
| GFP^+^ & D1R^+^ | NA | 1.25 ± 0.48 |
| IR8a^+^ & A_2A_R^+^ | 55.25 ± 4.52 | NA |
| IR8a^+^ & D1R^+^ | NA | 1.75 ± 0.48 |
| GFP^+^ & IR8a^+^ & A_2A_R^+^ | 50.50 ± 4.17 | NA |
| GFP+ & IR8a+ & D1R+ | NA | 1.25 ± 0.48 |

Notes: Lenti-FLEX-EGFP/IR84a-2A-IR8a vector (4.36 × 10^12^ genome copies/ml) was injected into the striatum of the *Drd2*-Cre rats and the striatal sections were subjected to two triple immunohistochemistry visualizations for GFP and IR8a with A_2A_R or D1R. Data are presented as mean cell numbers ± SEM of four samples.
